# Insights into the Polyhexamethylene Biguanide (PHMB) Mechanism of Action on Bacterial Membrane and DNA: A Molecular Dynamics Study

**DOI:** 10.1101/2020.03.25.007732

**Authors:** Shahin Sowlati-Hashjin, Paola Carbone, Mikko Karttunen

## Abstract

Polyhexamethylene biguanide (PHMB) is a cationic polymer with antimicrobial and antiviral properties. It has been commonly accepted that the antimicrobial activity is due the ability of PHMB to perforate the bacterial phospholipid membrane leading ultimately to its death. In this study we show by the means of atomistic molecular dynamics (MD) simulations that while the PHMB molecules attach to the surface of the phospholipid bilayer and partially penetrate it, they do not cause any pore formation at least within the microsecond simulation times. The polymers initially adsorb onto the membrane surface via the favourable electrostatic interactions between the phospholipid headgroups and the biguanide groups, and then partially penetrate the membrane slightly disrupting its structure. This, however, does not lead to the formation of any pores. The microsecond-scale simulations reveal that it is unlikely for PHMB to spontaneously pass through the phospholipid membrane. Our findings suggest that PHMB translocation across the bilayer may take place through binding to the phospholipids. Once inside the cell, the polymer can effectively ‘bind’ to DNA through extensive interactions with DNA phosphate backbone, which can potentially block the DNA replication process or activate DNA repair pathways.

**TOC Graphic:** 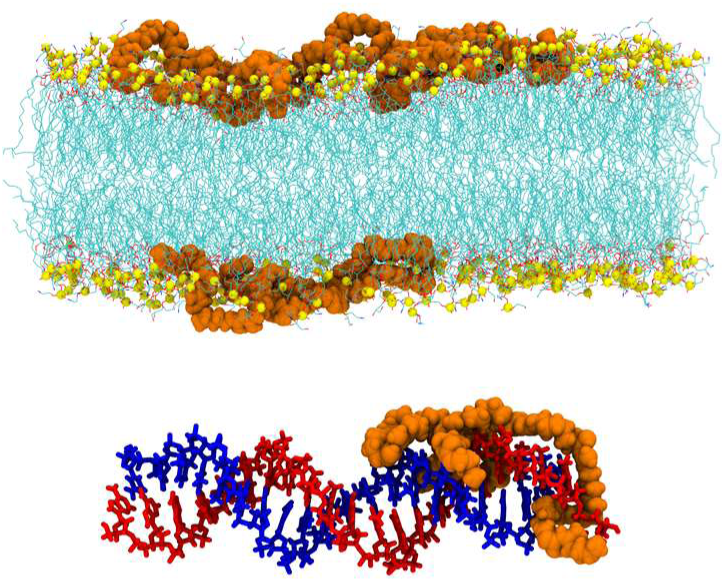

## Introduction

The cells of most of living organisms and viruses are separated from the surrounding environment by polar membranes that are made of lipid bilayers of different compositions. Similarly, cellular organelles in eukaryotes have their own characteristic lipid membranes that are important in determining their functionality.^1^ The lipid composition of membranes may vary from one species to another, yet its role in defining and protecting the cell, as well as controlling various physiological processes remains preserved in all organisms.^2^ Every exogenous factor (oxygen, ions, proteins, pathogens, etc.) first needs to pass through the cell membrane, and thus bacterial lipid membranes are a promising target for treatment of diseases. The matter is complex but, roughly speaking, the approaches are to either disrupt the membrane using antimicrobial molecules that damage the membrane or disturb its functions (such as permeability), or to target the enzymes that are responsible for bacteria’s ability to modify their membrane composition and thus develop resistance against antimicrobials.^3-4^

Biocidal cationic polymers such as polyhexamethylene biguanide (PHMB) are of great interest and widely used due to their high antibacterial and anti viral activity and low toxicity to humans. PHMB is mainly used as a disinfectant and antiseptic, and it has been employed, for example, in topical anti-infective solutions in ophthalmology,^5-6^ as disinfectants and biocides in water systems,^7^ and topically for wounds.^8^ It is also used as an ingredient in some contact lens cleaning products, cosmetics, personal deodorants, healthcare materials,^9^ and some veterinary products.^10-11^ Antimicrobial polymers such as PHMB have also shown to provide safe and efficient methods for gene delivery.^12-13^ In addition, PHMB doped silver nanoparticles (Ag NPs-PHMB) have been reported to reduce membrane fluidity, and cause cell membrane leakage.^14^ Strikingly, despite its antibacterial properties against various species^15-16^ and the fact that it has been used for decades, no bacterial resistance has been reported toward PHMB.^17^ Another fascinating feature of PHMB is its relatively low activity on the mammalian cell membranes.^18-22^ Other variants such as Polyhexamethylene guanidine hydrochloride (PHMG) have been studied and they have been shown to have marked impact on bacterial membrane functions.^23^

Despite numerous studies, PHMB mechanisms of action remain to be fully understood. Several studies have examined the interactions of PHMB with various membrane types concluding that PHMB readily interacts with negatively charged membranes^21, 24^ and that it is adsorbed on phospholipid bilayers.^25^ In addition to its interactions with membranes, PHMB has also been shown to induce DNA repair pathways.^26^ This observation cannot be solely explained by PHMB-membrane interactions. It has also been recently proposed that PHMB can enter both bacterial and mammalian cells and that it selectively condenses bacterial chromosomes.^27^ This contradicts the commonly accepted PHMB mechanism of action, in which the polymer disrupts the bacterial membrane.^18-19, 21^ According to this new proposal, a PHMB chain may enter a mammalian cell, but is unable to penetrate the nucleus where genetic material is stored. Since bacteria are prokaryotes and do not have membrane-bound organelles and nucleus, PHMB can therefore potentially directly interact with chromosomes upon entring the cell.

Computational studies of PHMB are currently limited to the study of structure of aqueous guanidinium chloride solutions,^28^ self-assembly of PHMB at various salt concentrations,^29^ adsorption on hydrogen peroxide treated Ti-Al-V alloys,^30^ and sorption on cellulose.^31^ At high concentrations, PHMB molecules aggregate and form mono- and multilayers on the surface through electrostatic interactions with counterions and hydrogen bonding of biguanide groups.

In this study, the dynamics and interactions of PHMB molecules in the form of dimer (containing two biguanide groups; Bgd^+^) and dodecamer (12 Bgd^+^ groups) in water solution with potassium chloride and on model bacterial phospholipid bilayers were studied to elucidate the effects of PHMB on them and to investigate the potential membrane penetration mechanisms. To further evaluate the PHMB-bilayer systems, and to examine the nature of the prevalent interactions and the possibility of salt bridge formation at the membrane interface, we considered the possible chemical reactions between PHMB and the reactive regions of the phospholipids (*i.e*., headgroups and unsaturated C=C bonds of the acyl chains). Finally, PHMB dodecamer was also allowed to interact with a DNA oligomer to study the PHMB-DNA contacts and the effects of such polymers on the structure and conformation of DNA.

## Computational Details

### Molecular Dynamics Simulations

The general protocol as described below was applied to all systems and system-specific details are given with the respective descriptions: After initial energy minimization using the steepest descents method and pre-equilibration (100 ps under constant volume and temperature (NVT) and 1 ns under constant pressure and temperature (NPT) for the PHMB-solvent systems, and 500 ps under NVT followed by 1 ns using NPT for the PHMB-bilayer systems), the production runs were carried out for at least 1 µs with 2 ps time step under NPT conditions with GROMACS/2018.^32^ Bonds were constrained with the linear constraint solver (P-LINCS).^33^ Temperature was maintained at 300 K using the Nosé-Hoover thermostat^34-35^ with a coupling constant of 1.0 ps. The Parrinello–Rahman barostat^36^ with a compressibility of 4.5 × 10^−5^ bar^−1^ and coupling constant of 5 ps was employed to keep the pressure constant. The long-range electrostatic interactions were evaluated using the particle-mesh Ewald method.^37-38^ Both the van der Waals and the real space electrostatic cut-offs were set to 1.2 nm. The OPLS-AA force field was used for PHMB and lipids parameters^29^ and the compatible TIP3P^39^ model for water.

### PHMB dimer and dodecamer chains in solvent

Chains of two different lengths that include the PHMB building block (**Figure S1a**) were examined: PHMB dimers consisting of two biguanide groups (**Figure S1b**) and PHMB dodecamer consisting of 12 biguanide groups (**Figure S1c**). For simplicity, from now on they are called dimer and polymer, respectively. In the present work, three systems were considered: single polymer, four polymers and four dimers in an explicit solvent. All systems were neutralized using Cl^−^ counterions and the whole system was then solvated in a water box with lengths of at least 1 nm from the edges of the molecules. The PHMB chains (both dimer and polymer) structures and parameters were obtained from Ref. 29.

### Bilayer and PHMB systems

The inner (cytoplasmic) membranes of Gram-negative bacteria, for example of *Escherichia coli*, characteristically consist of two main phospholipid types, phosphatidylethanolamine (PE, ∼70–80%) and phosphatidylglycerol (PG, ∼20–25%).^40^ The outer membrane, however, is mainly composed of lipopolysaccharide (LPS).^41^ Previous MD studies have modeled the inner membrane, examined various properties of such bilayers and revealed that many of them are controlled by the electrostatic interactions of the functional groups of the lipids with each other, ions and solvent.^42-44^ Recent simulations have also membrane proteins and the LPS.^45-47^ It has been shown that fully hydrated mixed lipid charged bilayers made of phosphatidylethanolamine (POPE) and phosphatidylglycerol (POPG) in the proportion 3:1 and counterions are stable and can serve as a suitable model of the inner bacterial membrane.^48^

In the present study, bacterial membranes were built using the CHARMM-GUI online server^49^ in two sizes: 1) To study PHMB polymer-bilayer systems, the bilayer consisted of 192 POPE and 64 POPG lipids (structures are provided in **Figure S2**) in each leaflet (total of 512 lipids), and 2) the bilayer for the PHMB dimer-bilayer system had 48 POPE and 16 POPG lipids in each leaflet (total of 128 lipids). A system of phospholipid bilayer similar to dimer-bilayer system was also prepared without PHMB chains to compare the bilayer properties in the absence of PHMB. Overall charge neutrality was preserved by adding counterions after which 0.15 M KCl was added. In total, the larger system had 217 K^+^ and 137 Cl^−^ ions and the smaller system 37 K^+^ and 13 Cl^−^ ions. The systems were solvated using a water box of 50708 and 5713 TIP3P^39^ water molecules in the two systems, respectively. The production phase for the polymer-bilayer system was 1 µs and the smaller dimer-bilayer system was simulated for 3 µs.

### PHMB–DNA System

Similar protocols as described above were used to set up the PHMB–DNA system. The DNA oligomer structure were taken from a topoisomerase I/DNA complex (PDB ID: 1A36,^50^ sequence (5′–AAAAAGACTTAGAAAAATTTTT)).

### Reactions

In order to evaluate the possibility of chemical reactions occurring between the polymer and phospholipids, the reaction potential energy surfaces (PESs) were investigated. First, transition structure corresponding to each reaction was characterized. Subsequently, the intrinsic reaction coordinate (IRC) was followed in both forward and reverse directions to obtain the product and reactant structures, respectively. The resulting structures were then fully optimized to obtain the relaxed reactants and products. This approach has been employed in numerous studies in both non-biological biological contexts, for example for studies of DNA damage and repair,^51-52^ and enzymatic catalysis.^53^ It is known to lead to reliable structures provided that the IRC is carefully scanned. All stationary points were fully optimized with B3LYP^54-56^ (B3LYP stands for Becke, 3-parameter, Lee-Yang-Parr) hybrid functional in combination with 6-311+G(2df,2p) level of theory in the presence of bulk solvent (water with ε = 78.3) as described by the IEF-PCM (Integral Equation Formalism-Polarizable Continuum Model) method^57-58^ implemented in Gaussian 16, Revision C.01.^59^

## Results and Discussion

### 1. PHMB in solvent

#### 1.1. Single PHMB chain (dodecamer) in solvent

Starting from a fully folded structure obtained from previous simulations at high salt concentrations,^12^ the PHMB polymer chain unfolded to some extent during the simulation but self-interactions of the chain were not fully lost (**Figure 1**). Root-mean-square-deviation (RMSD) indicates a drift from the initial folded conformation (**Figure S3**). This partial unfolding exposes the polymer to solvent and allows for creation of more hydrogen bonds with solvent. This leads to simultaneous reduction of self-interactions along the chain. There were on average 40 hydrogen bonds (H-bonds; (d(H⋯O) ≤ 0.2 nm and *α* (∠(Hydrogen–Donor–Acceptor) ≤ 30°) between the chain and solvent (**Table S1** and **Figure S4**), while the PHMB polymer formed only 1 hydrogen bond with itself. The average number of H-bonds per biguanide unit (Bgd^+^, **Figure S1a**) was found to be 3.4 (±0.3), which is slightly smaller, yet comparable to what was observed in high salt concentrations (4.0–4.5±0.6).^29^ Thus, self-interactions are mainly of van der Waals type. For all the systems considered in this study, the average number of hydrogen bonds per solvent molecule was found to be ∼3.1–3.3 (**Table S2**), which is comparable to what has previously been reported for bulk water.^60-61^

**Figure 1.**
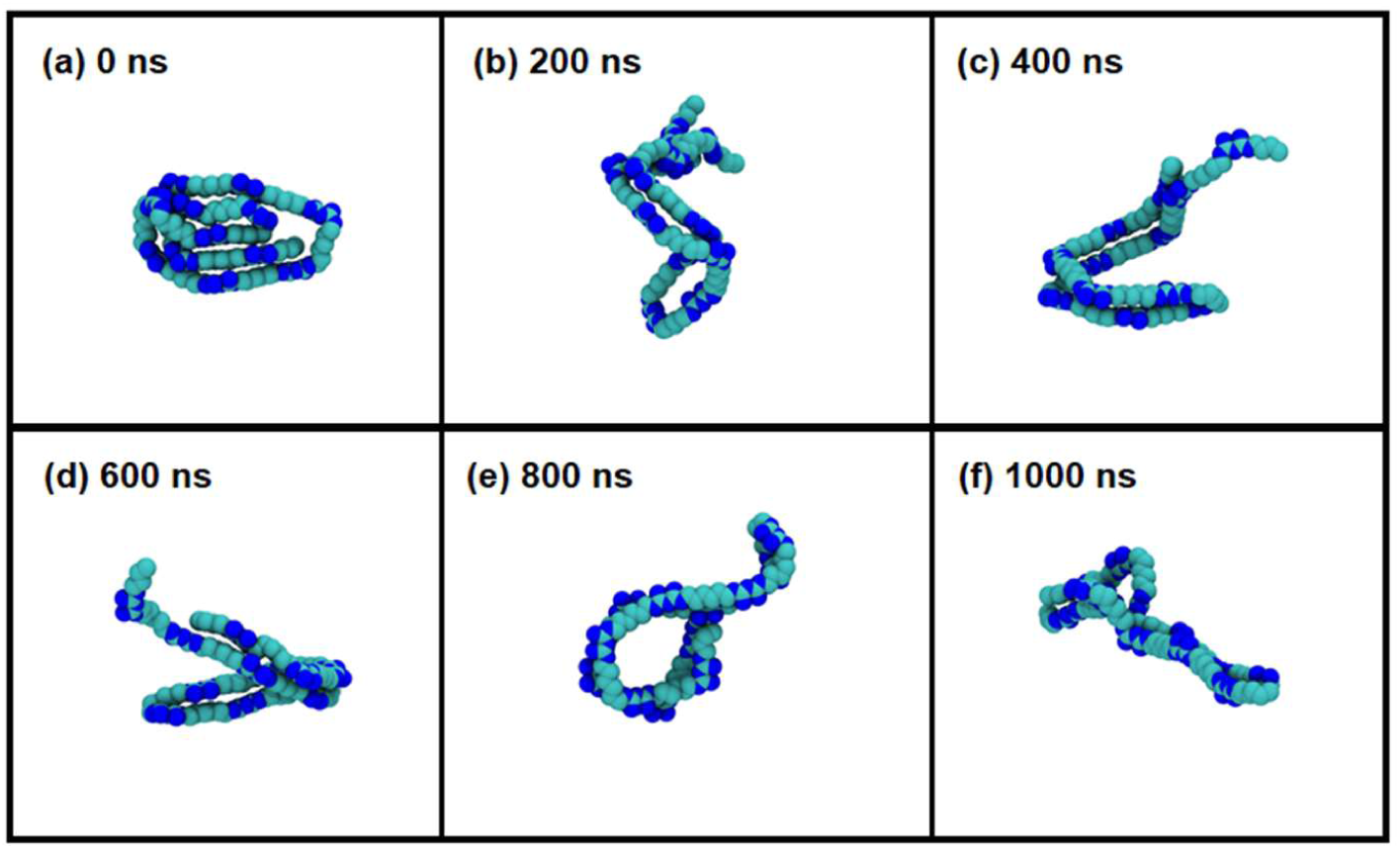
Snapshots of a single PHMB chain in solvent. Carbon and nitrogen atoms are shown in cyan and blue, respectively. Solvent molecules, hydrogen atoms and ions are not shown for clarity. Although the chain has lost its folded structure, completely unfolded structures were rare and the chain maintained its hairpin structure.

The polymer end-to-end distance is shown in **Figure S5a** and it reveals occasional complete unfolding of the chain. However, as is seen in **Figure 1**, the two ends of the chains remain close (1.57± 0.89 nm) as the polymer retains its semi-folded structure for the major part of the simulation. The polymer maintains its hairpin-like structure^29^ throughout the simulation, which keeps the polymer’s two ends close (**Figure 1**). Radius of gyration (R_g_) closely correlates with the end-to-end distance (**Figure S5b**). An incrsease/decrease in the distance means greater/smaller degrees of unfolding and hence, larger/smaller radius of gyration. The average R_g_ of the polymer is 1.55 (±0.23 nm). The solvent accessible surface area (SASA)^62^ of the polymer is also related to the end-to-end distance and R_g_, showing an increase at the beginning of the simulation and fluctuates around a constant throughout the time (31.60±2.01 nm^2^, **Figure S5c**).

#### 1.2. Four PHMB chains (dodecamer) in solvent

**Figure 2** shows snapshots of four PHMB chains in water starting from the exact same fully folded structure. The chains were initially a distance of at least 0.5 nm apart to warrant that no interactions among them prior to the simulation; presence of other chains has an impact on folding of PHMB to some extent. Although for the major portion of the simulation time the chains remained semi-folded as was also the case for a single PHMB chain (**Figure 1**), in some instances they fully unfolded and interacted with the neighbouring chain(s) (**Figure 2d**). The PHMB-solvent interactions were computed separately for each PHMB chain. The results show that the presence of other polymer chains, in comparison with the single polymer system, has negligible impact on the total number of hydrogen bond interactions between the chains and water. On average, each chain formed 38.0–39.6 hydrogen bonds with solvent during the simulation (**Table S1**). Similarly, the number of intra-hydrogen bonds for PHMB chains is not affected by the neighboring chains and remains at ∼1.1 (**Table S1, Figure S6**). This suggests that van der Waals interactions are the main component that governs polymer behaviour (folding, unfolding, self-interaction, etc.) and in high concentrations allow them to aggregate. Similar to the polymer-solvent system, the average number of solvent-solvent hydrogen bond was found to be 3.3 (±0.1) (**Table S2**).

**Figure 2.**
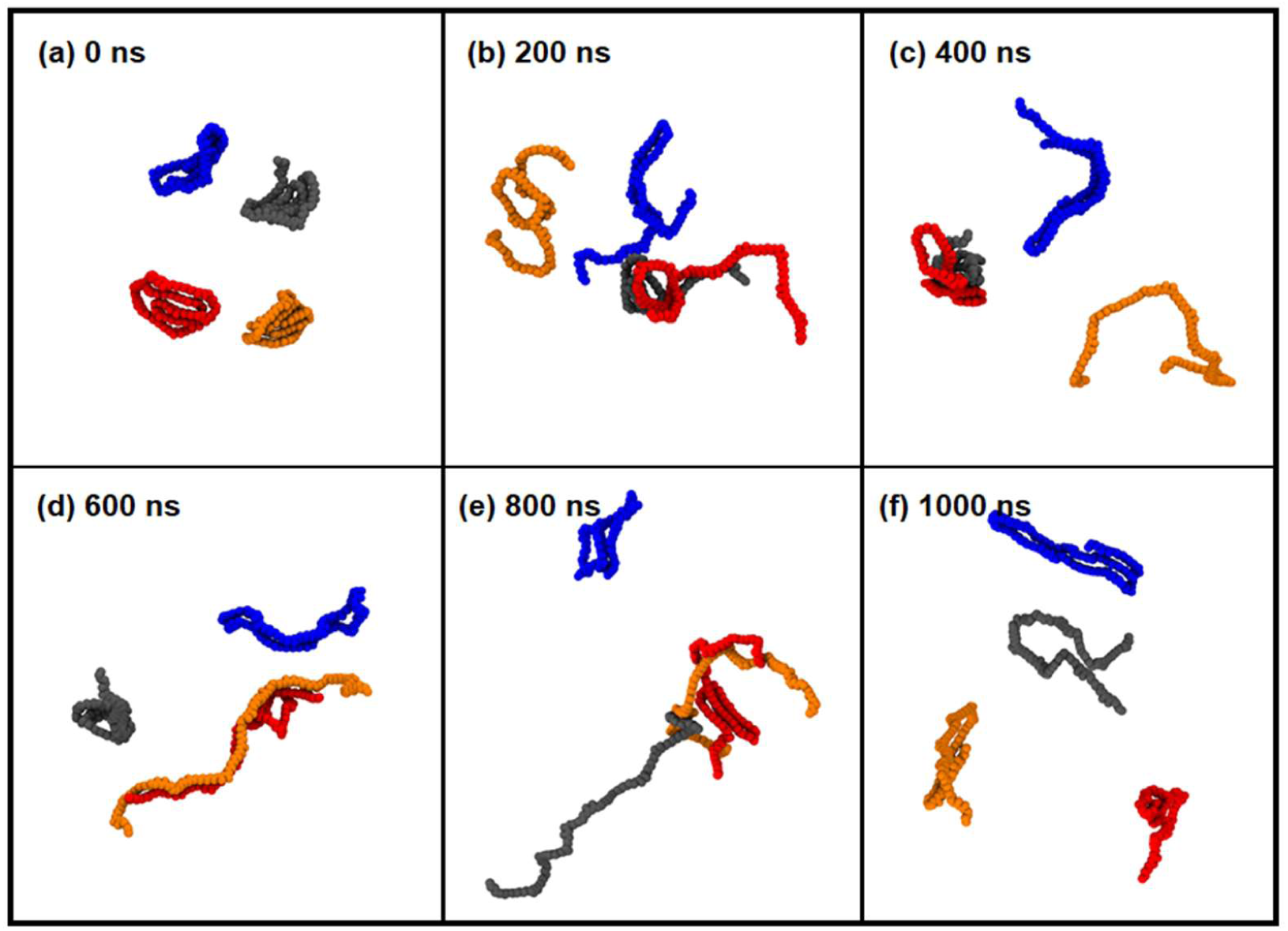
Snapshots of four PHMB chains in solvent. For clarity, the four chains are shown in different colors. Unlike in the case of one PHMB chain (Figure 1), there are frequent unfolding events and in some cases the hairpin structure unfolds. Solvent molecules, hydrogen atoms, and ions are not shown for clarity.

Compared to the single polymer system (1.57 nm), the average end-to-end distance of the polymer chains increased for the 4-polymer system (2.05, 4.22, 4.72, and 6.62 nm; **Figure S7**). In addition, as **Figure 2** shows, full unfolding occurs occasionally. Unfolding events lead to a larger average R_g_ (average of four chains: 2.18 nm, the largest ∼2.90 and the smallest ∼ 1.72 nm; **Figure S6**). The solvent accessible surface area (SASA) for the four chains shows larger fluctuations than what was observed for a single polymer (**Figure S7**). Interestingly, however, compared to the single polymer-solvent system (31.60±2.01 nm^2^), the average SASA for the four polymers is larger (35.30±3.23 nm^2^).

The end-to-end distance and R_g_ for the individual polymers are provided in **Figure S7**. The four polymers reveal larger average end-to-distance (4.40±1.52 nm) and R_g_ (2.18±0.29 nm) in comparison with the single polymer in solvent (distance=1.57±0.89 nm and R_g_=1.54±0.23 nm).The completely unfolded structures seen rarely in the single polymer-solvent system are observed more frequently when four chains are present (**Figure 2**).

#### 1.3. PHMB dimers in solvent

Compared to the PHMB polymer, the dimers showed a much lower propensity for mutual interaction and aggregation (**Figure 3**); the dimer chains only occasionally come close to each other. This observation is in clear contrast with a previous study^28^ and highlights the impact of Bgd^+^ concentration on the aggregation. This is discussed further in Conclusions. There are on average ∼6.9 H-bonds per chain and thus 3.5±0.8 H-bond per Bgd^+^ unit, which is in agreement with the number of H-bonds per unit in polymer systems (**Table S1** and **Figure S8**).

**Figure 3.**
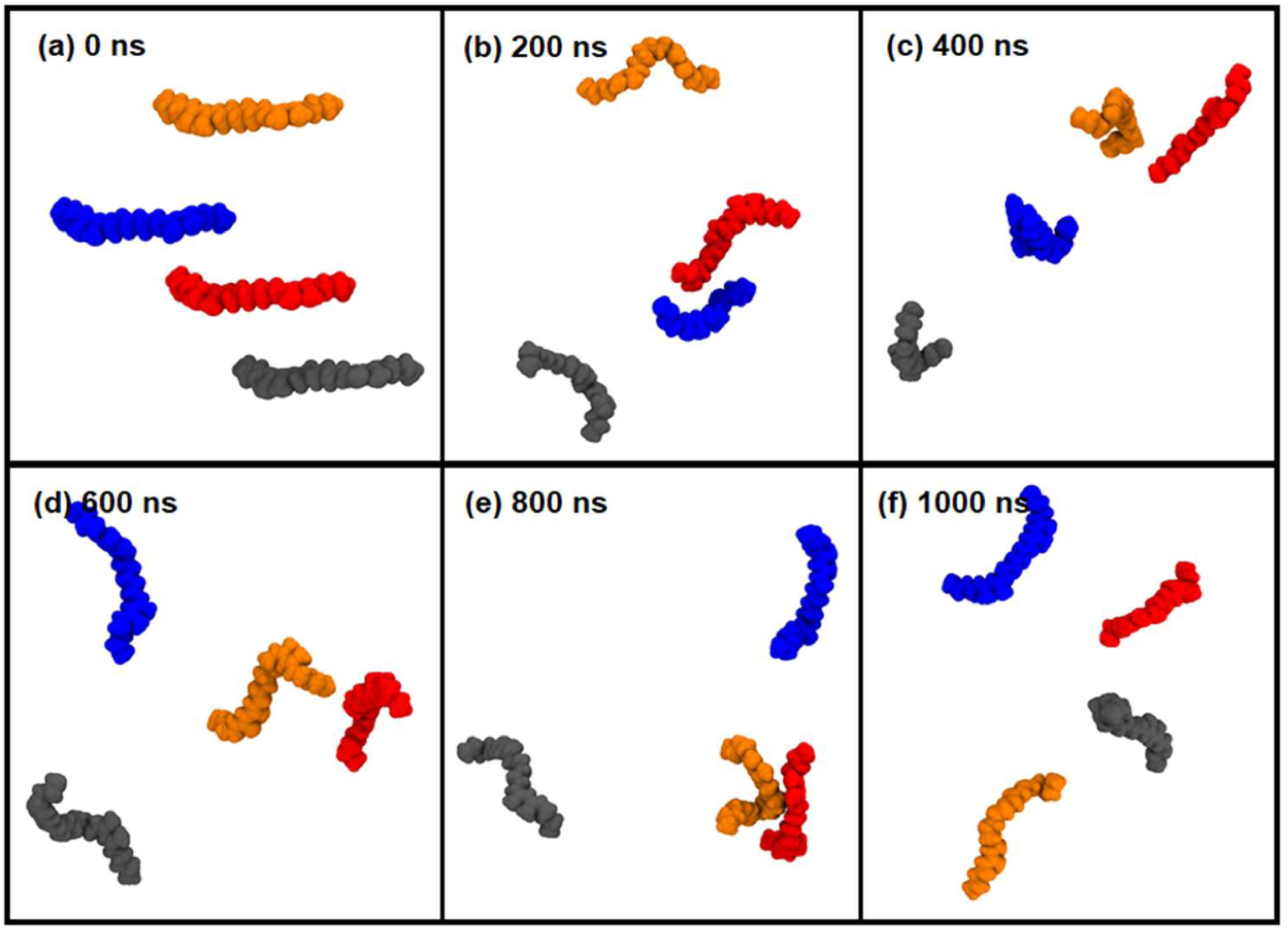
Snapshots of four dimers in solvent. The dimers do not show any tendency to fold or aggregate at this concentration (2.5 mM).

The short PHMB dimer chain is not as flexible as the longer PHMB polymer and hence for all four dimers, the end-to-end distance fluctuates between ∼0.5–2.5 nm (**Figure S9**). The radius of gyration (average ∼0.68 nm) closely follows the changes of the end-to-end distance with a small amplitude of fluctuation (∼0.4 nm; **Figure S9**). The average SASA for the four dimers (**Figure S9**) is 8.44 ± 0.31 nm^2^, which within the margin of error for SASA obtained for a single dimer in solvent 8.47 nm^2^, data not shown). This also indicates that there are negligible interactions among the dimer chains.

### 2. PHMB on bilayer

#### 2.1. Four PHMB chains (dodecamer) on bilayer

We studied systems with one PHMB and four PHMB (dodecamer) chains on a bilayer, but the discussion is limited to the latter. The four PHMB chains were placed in the solvent over the bilayer with a distance large enough to ensure that there were initially no direct interactions between the chains and the bilayer. Upon starting the simulation, the PHMB polymers immediately began to interact with the bilayer. **Figure 4** shows the relative positions of the PHMB chains on the bilayer during the simulation (corresponding snapshots along the bilayer normal are provided in **Figure S10**). Once on the bilayer, the motions of the polymers become restricted; the electrostatic interactions between the negatively charged phospholipids and positively charged chains as well as van der Waals interactions between the hydrophobic parts do not allow the polymers to move freely on the surface (**Figure 4**). Despite partial disruption of the bilayer by the polymers, consistent with previous reports on PHMB-phospholipid bilayer interactions,^21, 25^ no full penetration was observed.

**Figure 4.**
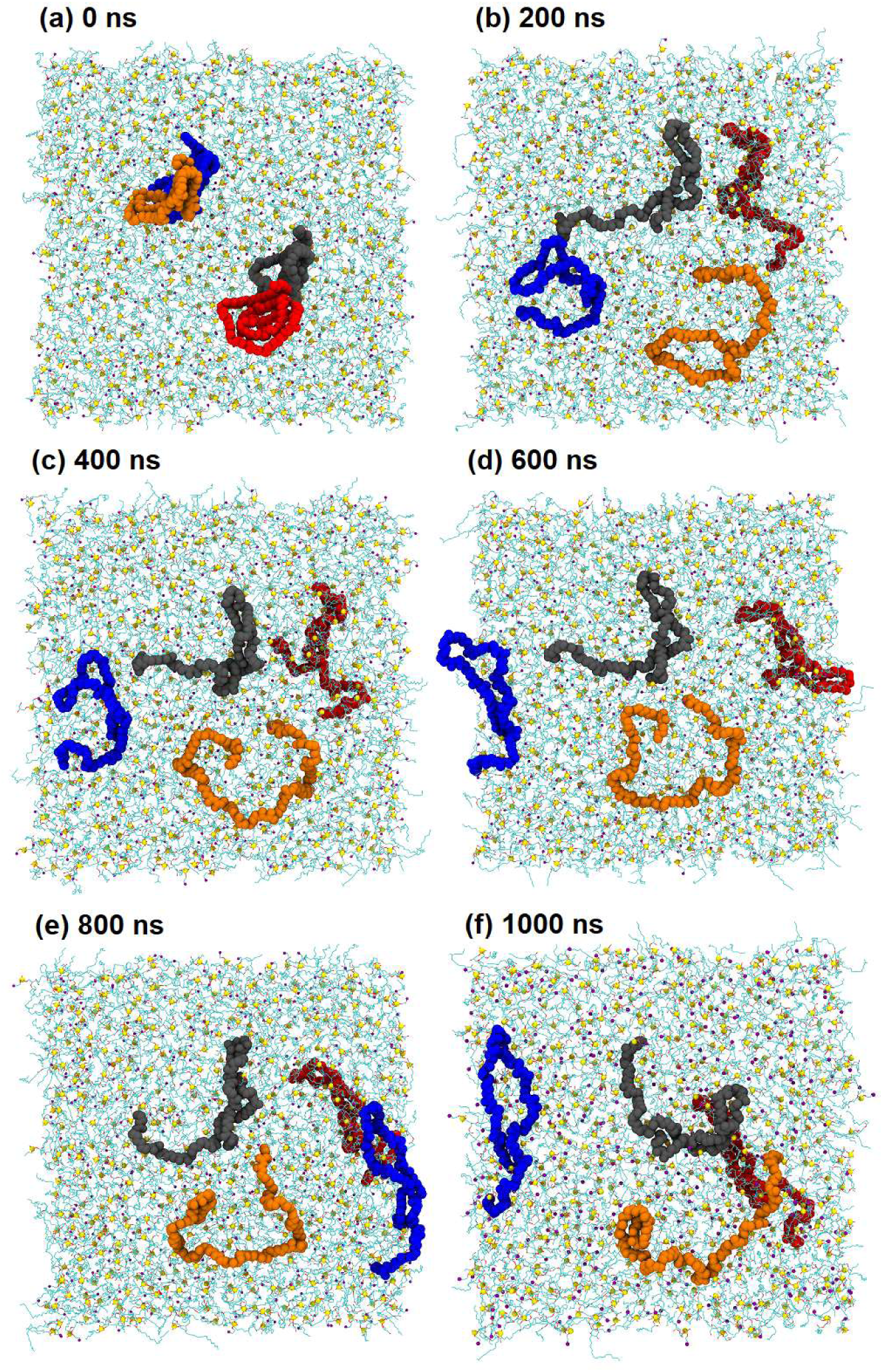
Snapshots of four polymers on the bilayer. Nitrogen is shown in purple, phosphorus in yellow and oxygen in red. Snapshots along the bilayer normal are provided in **Figure S11**. Once on the bilayer, the polymers are unfolded to some extent and their mobility is reduced.

When on a bilayer, the number of hydrogen bonds between the chains and solvent decreased significantly compared to the case of four polymers in water, ∼39 vs. ∼18 PHMB-solvent H-bonds, **Table S1**. Moreover, the average self-interaction decreased to < 1 hydrogen bond for the polymers on the bilayer (**Table S1** and **Figure S11**). This loss of hydrogen bonds was compensated by PHMB-bilayer interactions: ∼22 H-bonds per polymer chain were recorded, **Table S1**. Overall, compared to the four polymers in solvent, the number of hydrogen bonds formed per Bgd^+^ unit by the PHMB chains remains constant (∼3.5). The average numbers of POPG- and POPE-solvent hydrogen bonds were 5.8±0.1 and 5.8±0.1, respectively, that is, there was no significant change compared to the bilayer in the absence of PHMB (6.0 and 6.2; **Table S2**). These numbers are also comparable to those previously reported for PG (5.1) and PE (5.8) bilayers.^42^ It is also known that presence of counter ions and bridging waters can affect these numbers.^43^

The end-to-end distance does not experience large fluctuations (**Figure S11**). This is expected as the chains are more restricted compared to those that are in solvent (**Figure S7**). The average radius of gyration of the PHMB polymer chains on the bilayer (average of 4 chains: 1.57 ± 0.89 nm, the largest ∼1.78 and the smallest ∼1.44 nm; **Figure S12**) is reduced by ∼ 0.61 nm compared to the four PHMB polymer chains in solvent. Despite interactions with the bilayer, the average SASA of the PHMB polymers on the bilayer (35.01 ± 1.26 nm^2^; **Figure S12**) is essentially identical to the case of four PHMB polymers in a solvent (35.30 ± 3.23 nm^2^). This is likely due to the fact that on the bilayer the PHMB polymers have lost their folded structure and are stretched over the bilayer, which provides a larger surface area exposed to the solvent.

The density profile of the four polymers-bilayer system is shown in **Figure 5a**. As the figure shows, the PHMB polymer chains have partially penetrated the bilayer and PHMB density is the largest below the bilayer surface. This partial penetration can also be clearly seen in **Figure S10**, which shows snapshots of the PHMB-bilayer along the bilayer normal.

**Figure 5.**
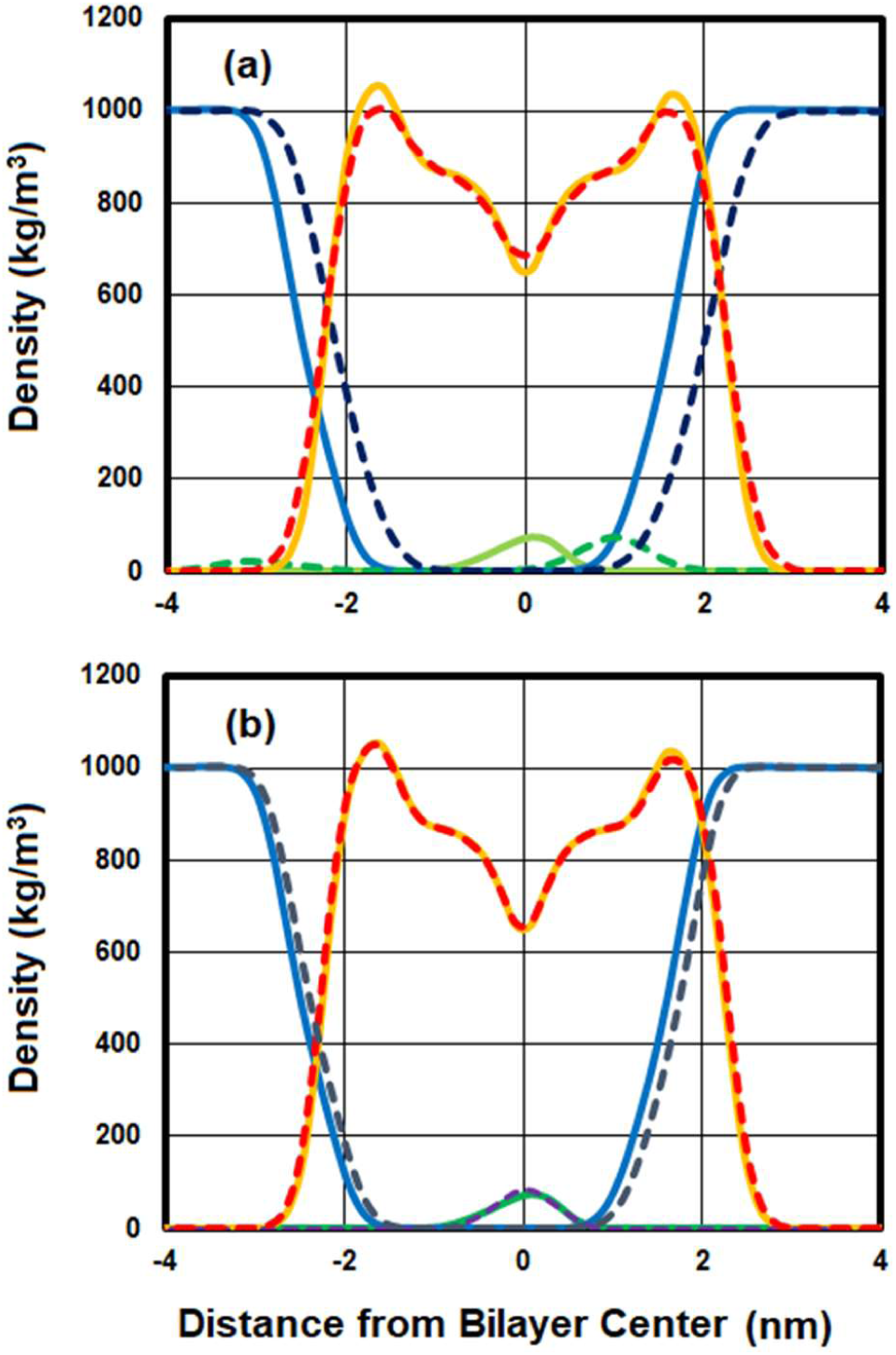
(a) Overlay of density profiles of four polymers-bilayer (dashed lines: red: bilayer, green: polymer, blue: water) and four dimers-bilayer (solid lines: orange: bilayer, green: dimer, blue: water). (b) Overlay of density profiles of our dimers-bilayer system at 1 µs (solid lines) and 3 µs (dashed red: bilayer, dashed dark blue: water, and dashed purple: dimers). As is seen, the dimers have penetrated the bilayer deeper compared to the polymers. This has resulted in unsymmetrical water distribution on the solvent-bilayer interface on the two sides of the bilayer (a). Density profile has not significantly changed after 1 µs in the presence of the dimers (b).

The area per lipid (APL) for the bilayer components (POPG and POPE) over time is shown in **Figure S13**. Average APLs for POPG are 0.63±0.01 nm^2^ and 0.62±0.03 nm^2^ and for POPE 0.57±0.01 nm^2^ and 0.56±0.02 nm^2^, for the PHMB polymer-bilayer and PHMB dimer-bilayer systems, respectively. The overall average APL of (0.61±0.01) nm^2^ for the PHMB polymer-bilayer does not undergo significant changes during the simulation (**Figure S13**). Interestingly, fluctuations in the APL for the PHMB dimer-bilayer system are significantly larger than for the polymer-bilayer system and the average APL is smaller (0.58 ± 0.01 nm^2^) compared to the PHMB polymer-bilayer system ((0.61±0.01) nm^2^). As a reference, we conducted an additional MD simulation of a bilayer system in the absence of PHMB chains for 100 ns. The average area per lipid in the absence of PHMB is smaller with the total value being 0.57±0.01 nm^2^ and component-wise POPG has 0.63±0.01 nm^2^ and POPE 0.56±0.01 nm^2^; the overall value of 0.57 ± 0.01 nm^2^ is in excellent agreement with a previous report on a PE/PG system (0.58±0.01 nm^2^).^63^ Although the different numbers of ions (*i.e*., K^+^ and Cl^−^) in the two systems (pure bilayer and PHMB-bilayer) may have some impact on the area per lipid,^64^ the effect of PHMB chains on the bilayer cannot be ignored. This is in line with previous data on expansion of PG bilayer (through insertion of hydrophobic hexamethylene groups into the non-polar region of the bilayer) at the water/air interface and increase of lateral distance between PG lipids in the presence of PHMB.^21^ The results indicate that polymer has an impact on the APL, while the effect of dimer is negligible. Larger average APL caused by the PHMB polymer may be indicative of greater exposure of the phosphate and carbonyl groups of the phospholipids. Bilayer thickness was also measured, and it follows the trends observed for APL as expected (**Figure S14**).

Order parameters of POPE and POPG acyl chains were evaluated separately over the first and last 100 nanoseconds of the simulations (*i.e*., 0–100 and 900–1000 ns) and are compared to those obtained for the bilayer in the absence of PHMB. The presence of PHMB induces slight but consistent disordering of both of the POPG chains. As the data show (**Figure S15a–b**), order decreases slowly with time. On the other hand, POPE order parameters appear not to be affected by the presence of PHMB (likely since there are more POPE lipids, and not all of them are involved, the effect of PHMB is not reflected in the average value). Concentrations of 2 – 6 ppm of PHMB has been previously shown to slightly increase the lipid order of DPPS-bilayer.^25^ The different bilayer composition, temperature, and PHMB concentration are the factors differentiating the previous experimental study^8^ and present results.

#### 2.2. Dimer chains on bilayer

Similar to the PHMB polymer chains, the PHMB dimers cannot move freely on the surface of the bilayer (**Figure 6**). The dimers penetrate the bilayer deeper than the longer polymers as can be seen in **Figures 5 and S16**. The average number of hydrogen bonds formed between the dimers and solvent is reduced to ∼2.3/dimer (**Table S1** and **Figure S17**) compared to the four dimer-solvent system (∼6.9 H-bonds/dimer; **Table S1** and **Figure S8**). This may be related to the fact that dimers penetrate the bilayer deeper and hence are less exposed to the solvent. On the other hand, the average number of H-bonds per Bgd^+^ with bilayer for the dimer-bilayer is larger than that of the polymer-bilayer system. The presence of the dimers does not seem to affect bilayer-solvent hydrogen bonding and there are on average 5.9 or 5.8 hydrogen bonds between the solvent and POPG or POPE, respectively (**Table S2**).

**Figure 6.**
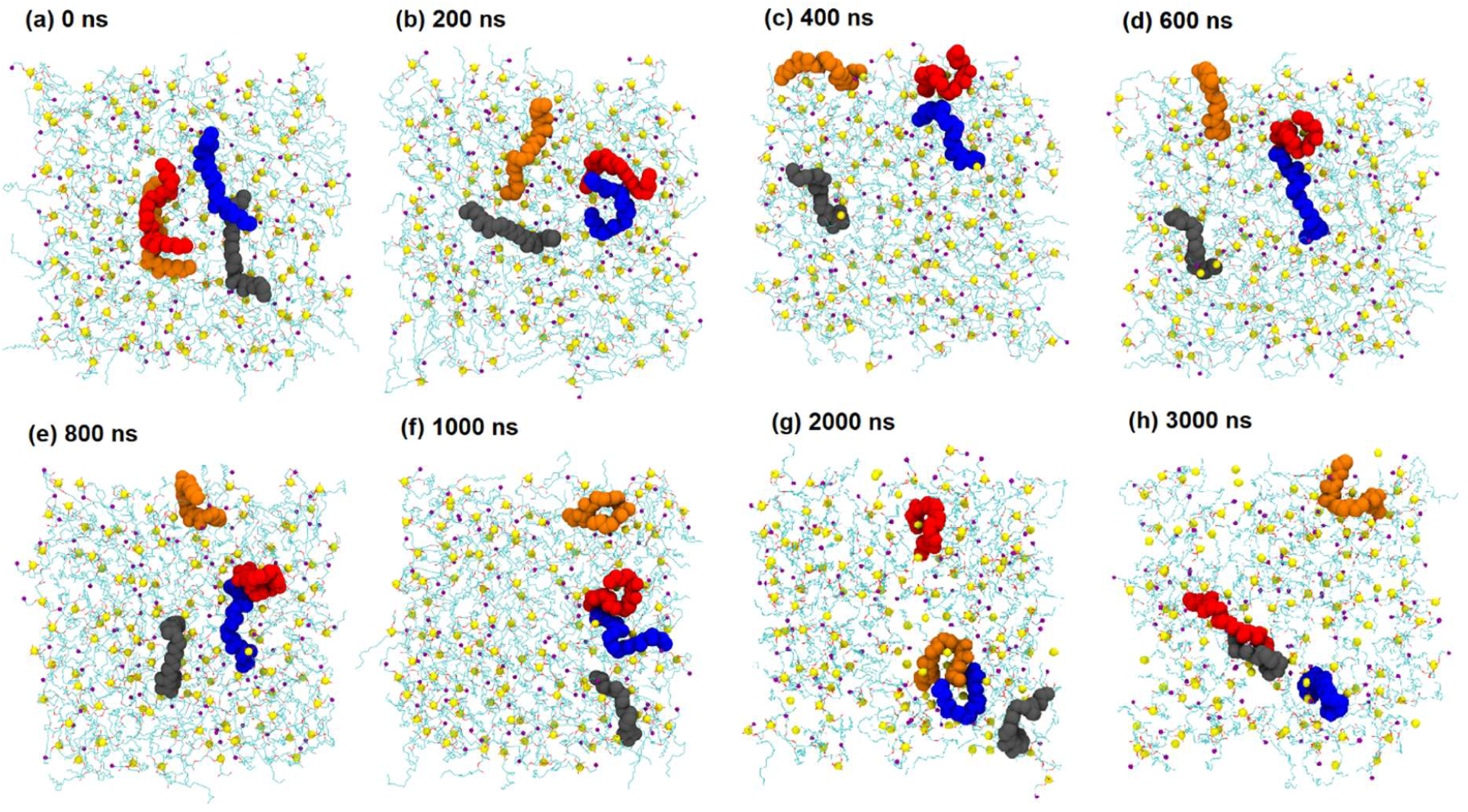
Snapshots of the relative positions of the four dimers on the bilayer during the simulation time. Nitrogen is shown in purple, phosphorus in yellow and oxygen in red. Snapshots along the bilayer normal are provided in **Figure S16**.

The average end-to-end distance (1.02±0.11 nm) and fluctuations are significantly reduced for the dimers on the bilayer (**Figure S18**) and the chains are mainly in their bent forms. The radius of gyration for the dimers is also decreased on the bilayer (0.55±0.06 nm; **Figure S18**). The SASA of the dimers in solvent (8.44±0.31 nm^2^) also decreases (8.06±0.37 nm^2^) in the dimer-bilayer system (**Figure S18**). This is in line with the reduction of number of H-bonds compared to the 4 dimers-solvent system.

The density profiles (**Figure 5**) reveal important differences between the dimer- and polymer-bilayer systems. For example, in comparison with the polymer, the dimers are located deeper in the bilayer, exposing the deeper regions of the bilayer to the solvent. The average area per lipid of the bilayer in the presence of the dimers remains constant at 1 and 3 µs and is (0.58±0.01) nm^2^, marginally larger than APL for the pure bilayer (0.57±0.01 nm^2^, at 100 ns) and smaller (∼ 3 nm^2^) than that of the polymer-bilayer system (0.61±0.01 nm^2^ after 1 µs; **Figure S13**). Other than the sn-2 chain of POPG, which is slightly affected by the presence of the dimers, other order parameters are not affected by the dimers over time (**Figure S19**).

#### 2.3. Reactions

It is been commonly accepted that PHMB disrupts bacterial membrane.^18-21, 25^ It is not clear, however, if this disturbance is solely physical, taking place through chemical reactions or if it is a combination of the two. Moreover, it has been proposed that cationic-penetrating peptides may be transferred into the cell by forming salt bridges with phospholipid negatively charged headgroup.^65-68^ To resolve this issue, we examined all potential chemical reactions between the bilayer (phospholipids) and the biguanide group. It turned out, however, to be impossible to characterize all of them, since several of the transition structures could not be isolated. Those cases were discarded (**Figure S20**). Reactions that completed are shown in **Figure 7**. The high energy barriers suggest that the disruption of membrane by PHMB is unlikely and can be excluded. However, it is possible for covalent bonds to form between PHMB and lipids. Although chemical links between PHMB and the lipid are unlikely, the opposite Bgd^+^ and phosphate charges warrant formation of PHMB-lipid salt bridge and promote the translocation of PHMB.

**Figure 7.**
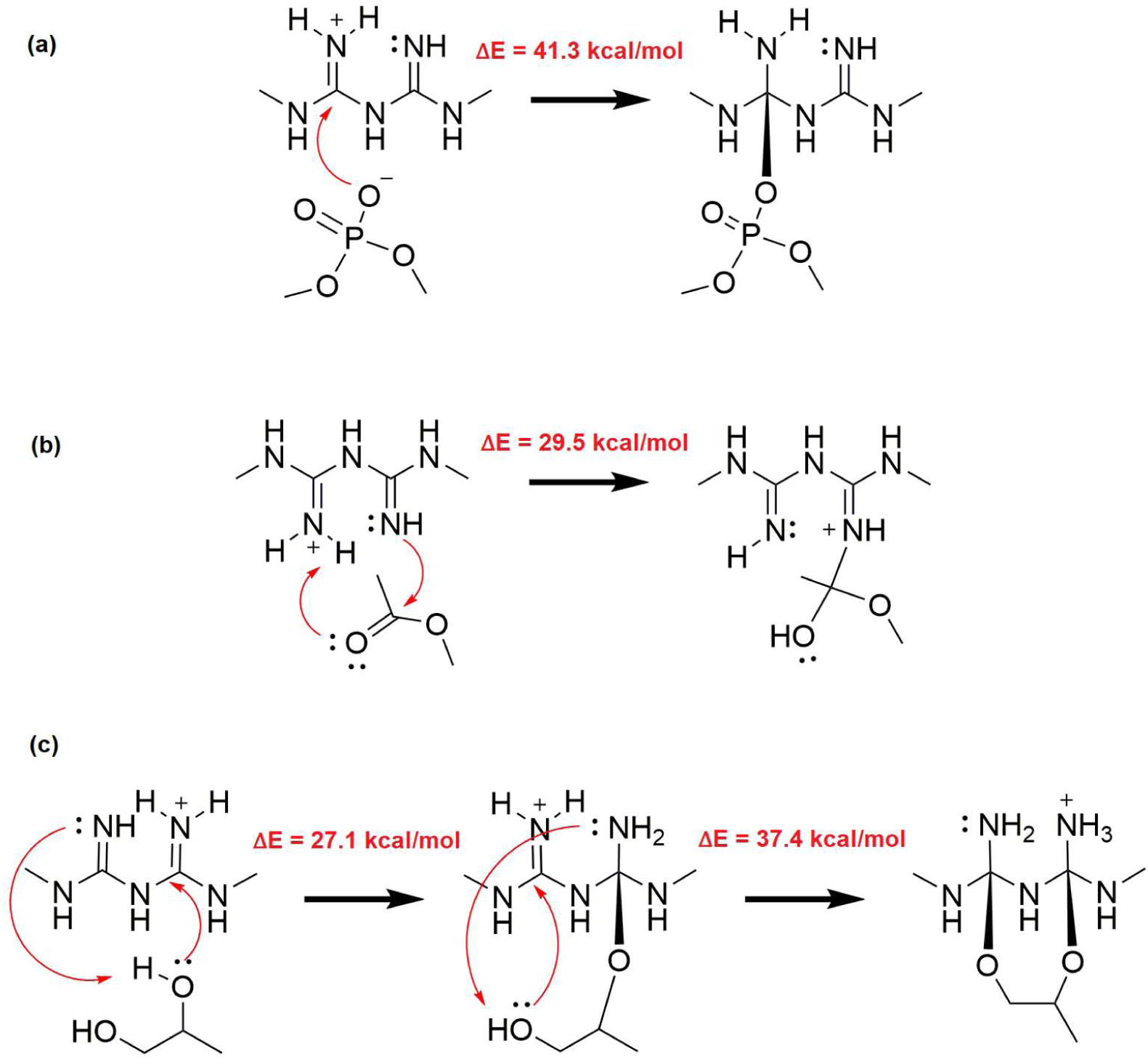
Reactions between the biguanide group and (a) phosphate, (b) the POPG head group, and (c) the ester group of the phospholipid chains. Energy barriers for the reactions suggest the unlikeliness of such reactions.

### 3. PHMB–DNA interactions

In addition to membrane damage, it has been proposed that PHMB may penetrate through the membrane and activate DNA repair pathways.^26^ Our MD simulations reveal extensive interactions between the PHMB chain and DNA (**Figure 8**). Specifically, the number of PHMB– DNA hydrogen bonds rapidly increases from zero at the beginning of the simulation (**Figure 8a**) to an average of 21 (**Table S3 and Figure S21**). Meanwhile the number of H-bonds between the PHMB chain and solvent decreases. The average number of total DNA base-paring hydrogen bonds remains constant at 42±2, which is only three hydrogen bonds short of its ideal number according to the DNA oligomer sequence (5′–AAAAAGACTTAGAAAAATTTTT). This preserved hydrogen bonding indicates that major PHMB and DNA interactions take place at the DNA backbone (*i.e.*, phosphate groups).

**Figure 8.**
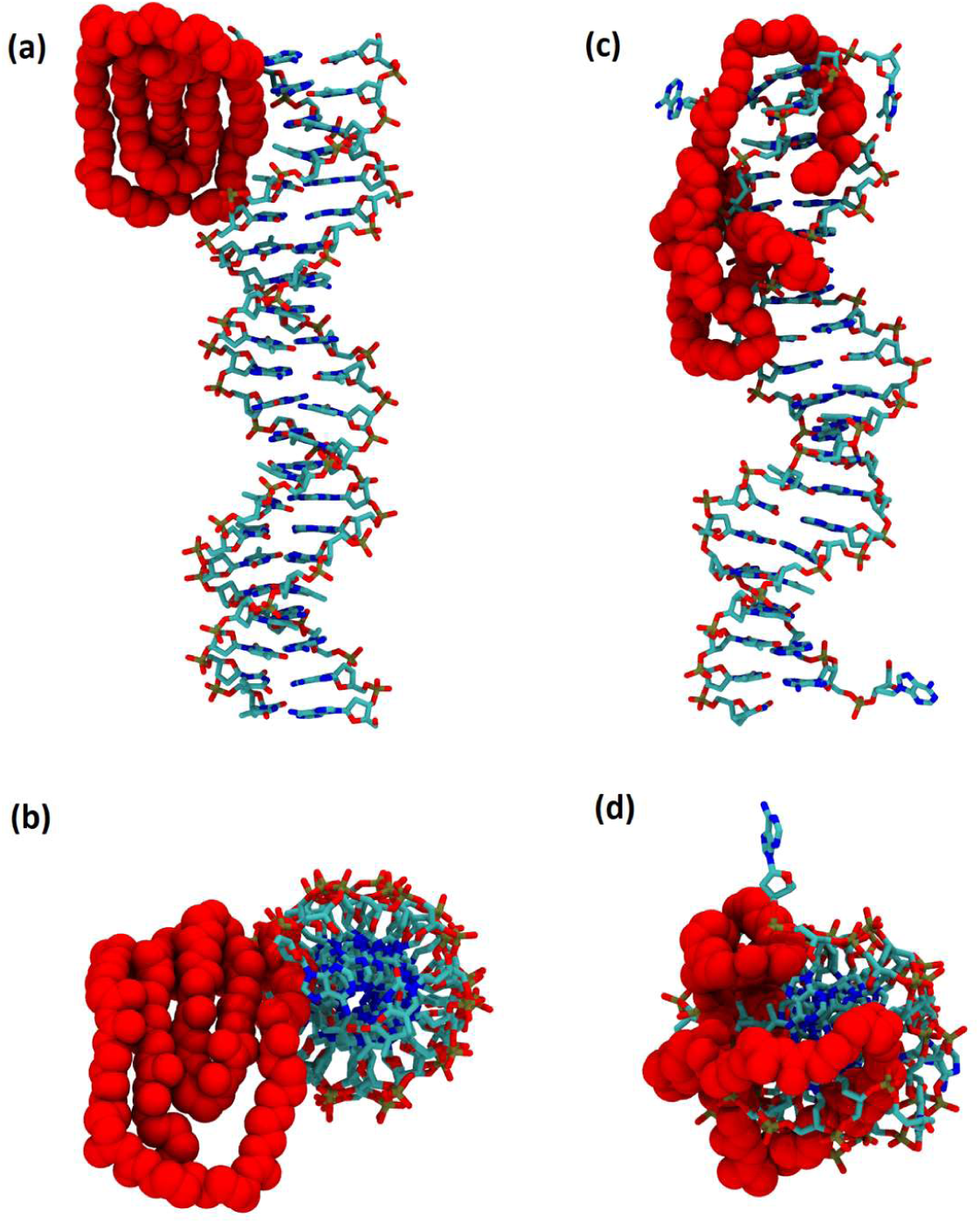
Initial (a: side-view and b: top-view) and final (1 µs; c: side-view and d: top-view) structures of the PHMB–DNA system. As seen, PHMB–DNA interactions (H-bond) increase with time.

During the early stages of the simulation (∼ 50–100 ns) a nucleobase pair dislocation was observed (**Figure S22**); however, this anomaly did not last, and the base pair was retrieved. The position of the base-pair dislocation relative to the PHMB polymer suggests that the PHMB chain impacts on the DNA conformation mainly through interactions with the backbone. As is seen in **Figure 8**, the PHMB chain remains folded to a great extent despite being in direct contact with a DNA. This is also reflected in the RMSD and radius of gyration (**Figure S23**). Despite slight penetration and interactions at the major and minor grooves, the PHMB chain remains in the vicinity of the phosphate backbone. These interactions seem to disturb the overall symmetry of the DNA (**Figure 8d**). Moreover, tight binding of the PHMB chain to the DNA may block the DNA replication or activate DNA repair pathways. This is in line with a recent study on gene condensation through binding of PHMB to DNA^13^ and effective inhibition of DNA replication by Ag NPs-PHMB.^14^

## Conclusions

In the present study, in order to gain insight on the PHMB mechanism of action on the bacterial membrane, we examined the dynamic properties of PHMB dimer and dodecamer chains and their interactions with phospholipid bilayers.

It has been shown that PHMB units stack and form aggregates at high concentrations.^28^ Our MD simulation results suggest that at low concentrations of PHMB, the PHMB chains fold in a way so that biguanide units stack on top of each other. Our microsecond simulations on PHMB chains in solvent is consistent with the previous study on the self-assembly of the polymer in various high salt concentration.^29^ This stacking behaviour is seen for inter- and intra-chain interaction between PHMB units and is more evident at high salt concentrations.^29^ More specifically, at zero salt concentrations PHMB dodecamers partially unfold; however, stacking of PHMB units is never completely lost.

Previous experimental data suggest that PHMB only interacts with negatively charged phospholipid membranes and only above critical concentrations adsorption of PHMB can significantly reduce the bilayer stability.^21^ The results on the PHMB-bilayer systems (with low concentrations of dimers or dodecamers) indicate that spontaneous full penetration of the chain is unlikely to occur in the scale of 1–3 µs. However, penetration may occur through mechanisms proposed for cationic penetrating peptides.^65-68^ Specifically, formation of salt bridges between the cationic PHMB and anionic phospholipids may facilitate the translocation of the former across the bilayer. The hypothesis that PHMB mechanism of translocation may occur through binding is also supported by the observed extensive interactions between the opposite charges on the PHMB chains and the phospholipid bilayer, as well as hydrogen bonds between PHMB and lipids. The process may, however, be slow in terms of molecular simulations and it may be assisted by the compositional differences between the membrane leaflets. Intensive PHMB-bilayer interactions on the surface of different bilayers has been reported^21, 25^ and strengthen the possibility of such a hypothesis. Furthermore, the feasibility of bilayer disruption with potential chemical reactions between the functional group of the polymer and phospholipids can be ruled out due to the high energy barriers (∼ 27–41 kcal/mol), considering the number of reactions required to occur in order to have a large impact on the bilayer. Another possible mechanism that may apply to PHMB translocation, as proposed for transportan,^69^ is stochastic fluctuations of the PHMB chain toward the opposite monolayer of the membrane once it has reached to the center of bilayer. This is more probable for the PHMB dimers since they penetrate the bilayer deeper. Both proposed mechanisms, stochastic penetration and translocation facilitated by salt bridge formation, require intensive interactions between PHMB and the lipid bilayer, which is more pronounced for the anionic bacterial membranes.

Regardless of the mechanism by which PHMB may enter the prokaryote cells, it is shown that PHMB dodecamer extensively interacts with the DNA backbone and adopts a lump-like formation. This aggregate, specially when occurs with large number of PHMB chains, can potentially block the DNA replication pathway and/or induce DNA repair processes.

Overall, our findings suggest that disruption of membrane by PHMB is unlikely to be the dominant mechanism through which PHMB acts on bacterial membranes. The data presented in this study support the recent proposal^27^ on the translocation of PHMB across the bacterial membrane and acting directly on the genetic material. We hypothesise that translocation of PHMB probably takes place through a similar mechanism proposed for cell penetrating peptides, i.e., binding to the lipids.

## Acknowledgments

Financial support of this work was provided by the Natural Sciences and Engineering Research Council of Canada (NSERC). MK also thanks NSERC Canada Research Chairs Program. Computational resources were provided by Compute Canada.

## Supplementary Information

**Table S1.**
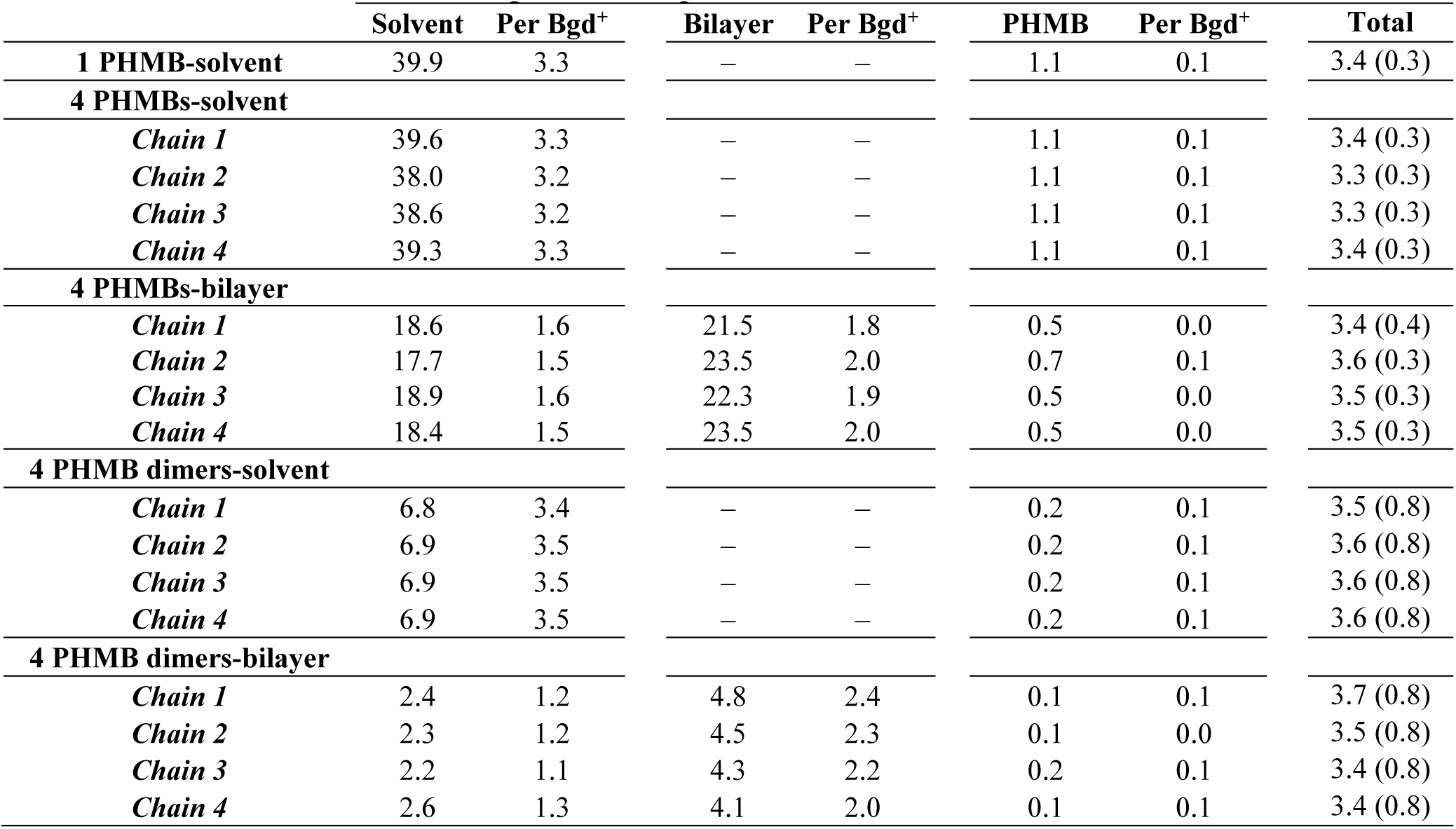
Average number of hydrogen bonds (d(H⋯O) ≤ 0.2 nm and *α* (∠(Hydrogen–Donor– Acceptor) ≤ 30°) for PHMB chains (with solvent, bilayer, and self interaction) in the different systems. Standard deviations are provided in parentheses.

|  | **Solvent** | **Per Bgd<sup>+</sup>** | **Bilayer** | **Per Bgd<sup>+</sup>** | **PHMB** | **Per Bgd<sup>+</sup>** | **Total** |
| --- | --- | --- | --- | --- | --- | --- | --- |
| **1 PHMB-solvent** | 39.9 | 3.3 | – | – | 1.1 | 0.1 | 3.4 (0.3) |
| **4 PHMBs-solvent** |  |  |  |  |  |  |  |
| *Chain 1* | 39.6 | 3.3 | – | – | 1.1 | 0.1 | 3.4 (0.3) |
| *Chain 2* | 38.0 | 3.2 | – | – | 1.1 | 0.1 | 3.3 (0.3) |
| *Chain 3* | 38.6 | 3.2 | – | – | 1.1 | 0.1 | 3.3 (0.3) |
| *Chain 4* | 39.3 | 3.3 | – | – | 1.1 | 0.1 | 3.4 (0.3) |
| **4 PHMBs-bilayer** |  |  |  |  |  |  |  |
| *Chain 1* | 18.6 | 1.6 | 21.5 | 1.8 | 0.5 | 0.0 | 3.4 (0.4) |
| *Chain 2* | 17.7 | 1.5 | 23.5 | 2.0 | 0.7 | 0.1 | 3.6 (0.3) |
| *Chain 3* | 18.9 | 1.6 | 22.3 | 1.9 | 0.5 | 0.0 | 3.5 (0.3) |
| *Chain 4* | 18.4 | 1.5 | 23.5 | 2.0 | 0.5 | 0.0 | 3.5 (0.3) |
| **4 PHMB dimers-solvent** |  |  |  |  |  |  |  |
| *Chain 1* | 6.8 | 3.4 | – | – | 0.2 | 0.1 | 3.5 (0.8) |
| *Chain 2* | 6.9 | 3.5 | – | – | 0.2 | 0.1 | 3.6 (0.8) |
| *Chain 3* | 6.9 | 3.5 | – | – | 0.2 | 0.1 | 3.6 (0.8) |
| *Chain 4* | 6.9 | 3.5 | – | – | 0.2 | 0.1 | 3.6 (0.8) |
| **4 PHMB dimers-bilayer** |  |  |  |  |  |  |  |
| *Chain 1* | 2.4 | 1.2 | 4.8 | 2.4 | 0.1 | 0.1 | 3.7 (0.8) |
| *Chain 2* | 2.3 | 1.2 | 4.5 | 2.3 | 0.1 | 0.0 | 3.5 (0.8) |
| *Chain 3* | 2.2 | 1.1 | 4.3 | 2.2 | 0.2 | 0.1 | 3.4 (0.8) |
| *Chain 4* | 2.6 | 1.3 | 4.1 | 2.0 | 0.1 | 0.1 | 3.4 (0.8) |

**Table S2.** Average number of phospholipid-solvent and solvent-solvent hydrogen bonds (d(H⋯O) ≤ 0.2 nm and *α* (∠(Hydrogen–Donor–Acceptor) ≤ 30°). Standard deviations are provided in parentheses.

|  | **POPE⋯solvent** | **POPG⋯solvent** | **solvent⋯solvent** |
| --- | --- | --- | --- |
| **1 PHMB-solvent** | – | – | 3.3 (0.1) |
| **4 PHMBs-solvent** | – | – | 3.3 (0.1) |
| **4 PHMBs-bilayer** | 5.8 (0.1) | 5.8 (0.2) | 3.3 (0.1) |
| **4 PHMB dimers-solvent** | – | – | 3.3 (0.1) |
| **4 PHMB dimers-bilayer** | 5.8 (0.2) | 5.9 (0.3) | 3.1 (0.1) |
| **Pure bilayer** | 6.0 (0.2) | 6.2 (0.3) | 3.1 (0.1) |
| **Pure bilayer\*** | 5.8 | 5.1 | – |

\*Data from reference 2

**Table S3.** Average number of hydrogen bonds (d(H⋯O) ≤ 0.2 nm and *α* (∠(Hydrogen–Donor– Acceptor) ≤ 30°) in the PHMB-DNA system. Standard deviations are provided in parentheses.

|  | **PHMB⋯solvent** | **Per Bgd<sup>+</sup>** | **Total** | **DNA⋯DNA** |
| --- | --- | --- | --- | --- |
| **1 PHMB-DNA** | 23.3 | 1.9 | 3.1 (0.3) | 42.9 (2.4) |
|  | **PHMB⋯DNA** | **Per Bgd<sup>+</sup>** |  |  |
|  | 13.0 | 1.1 |  |  |
|  | **PHMB⋯PHMB** | **Per Bgd<sup>+</sup>** |  |  |
|  | 1.0 | 0.1 |  |  |

**Figure S1.**
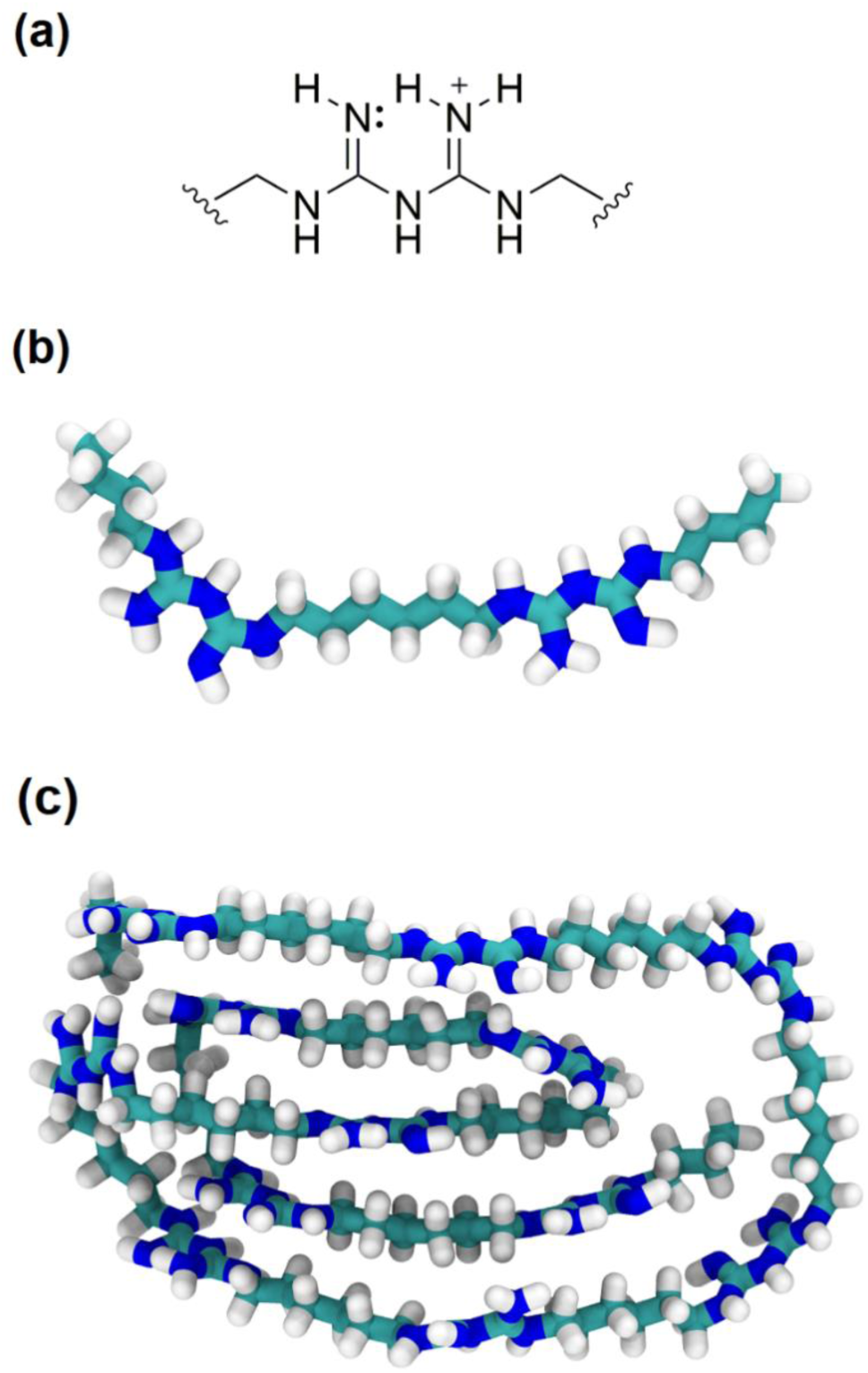
(a) 2D representation of the biguanide functional group. Structures of biguanide (b) dimer and (c) dodecamer. Carbon, nitrogen and hydrogen atoms are shown in cyan, blue and white, respectively.

**Figure S2.**
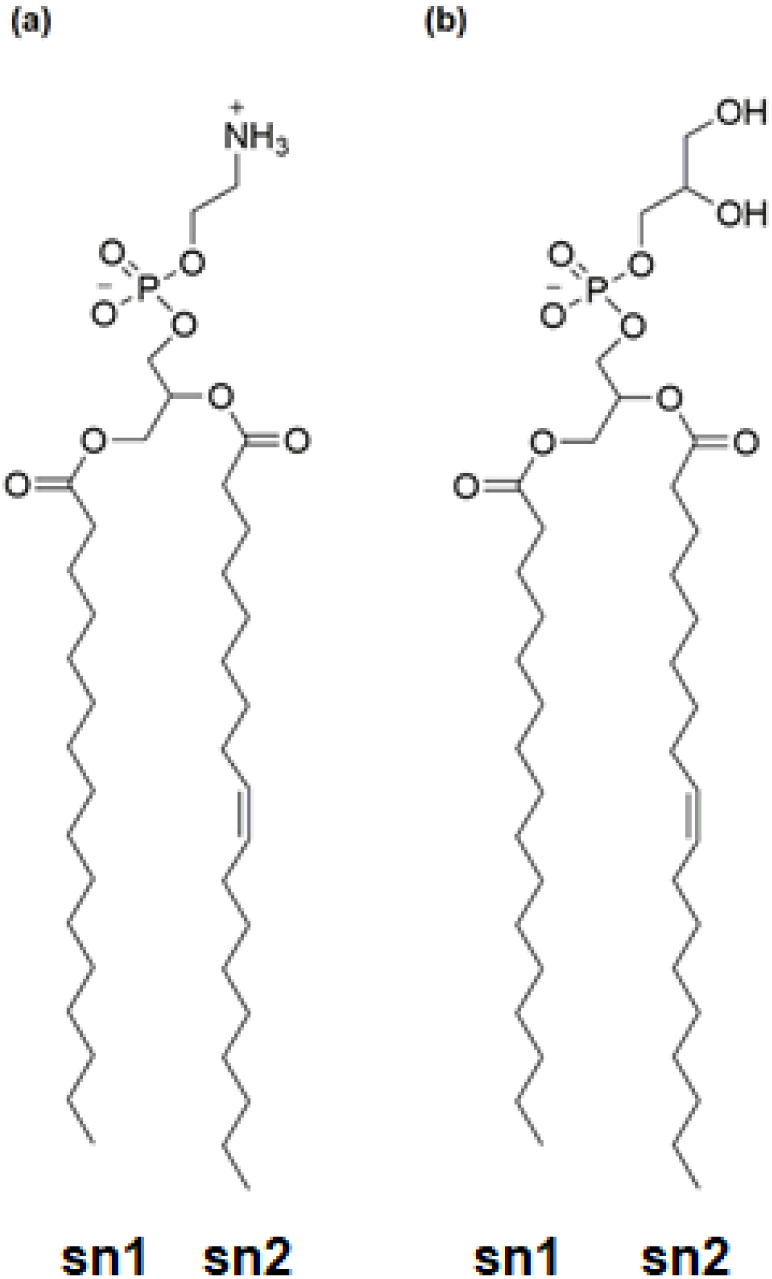
2D representations of (a) phosphatidylethanolamine (POPE) and (b) phosphatidylglycerol (POPG) lipids.

**Figure S3.**
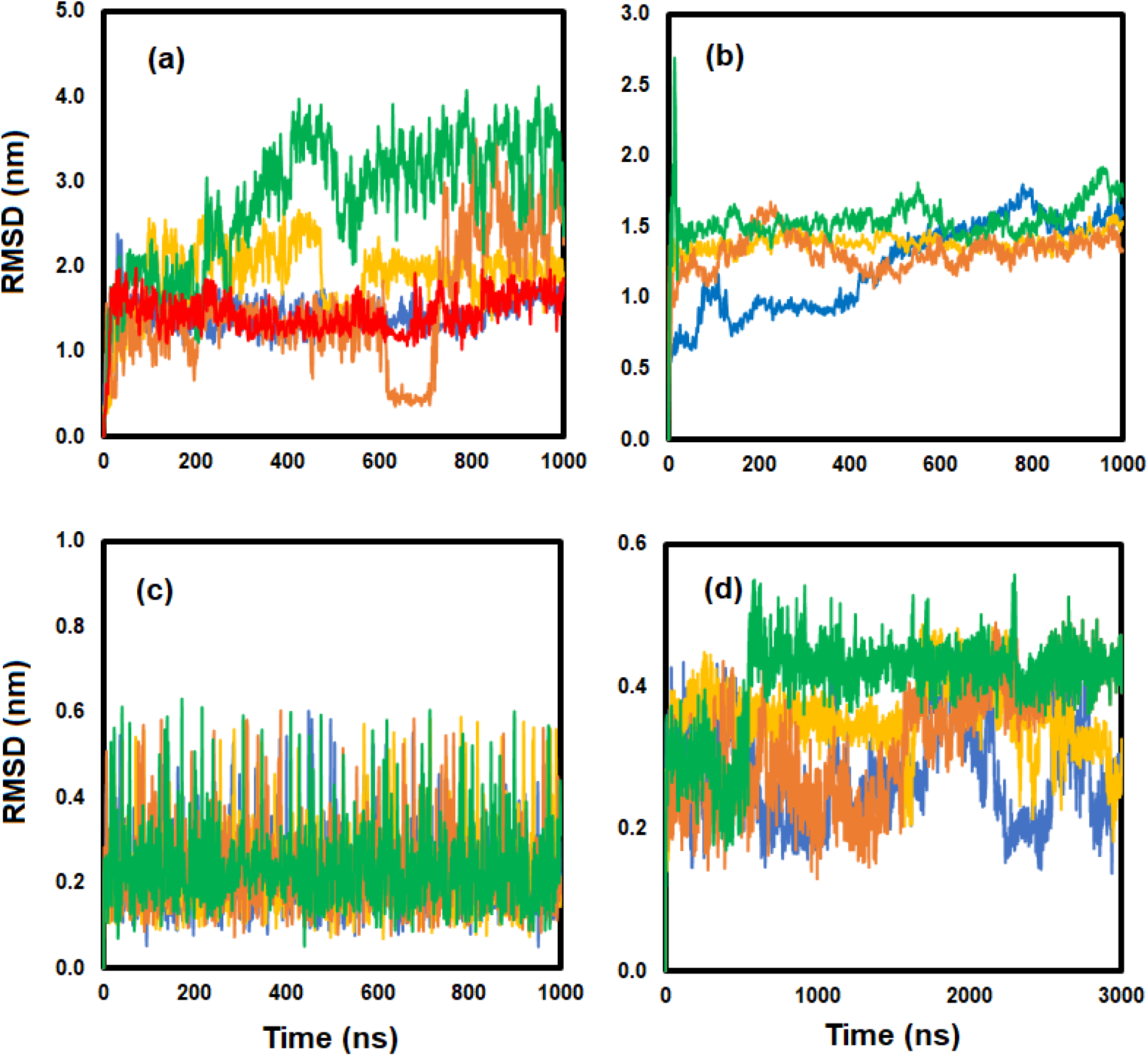
RMSD of (a) 1 PHMB polymer (red) and 4 PHMB polymers in solvent (individual chains are represented in different colors: green, blue, orange, and brown). (b) RMSD of 4 PHMB polymers on the bilayer. (c) RMSD of 4 PHMB dimers in solvent and (d) of 4 PHMB dimers on the bilayer. As is seen, in comparison with the solvent system, both the PHMB polymer and dimer chains show lower RMSDs on the bilayer, indicating a reduced degree of mobility when involved in interactions with the phospholipids.

**Figure S4.**
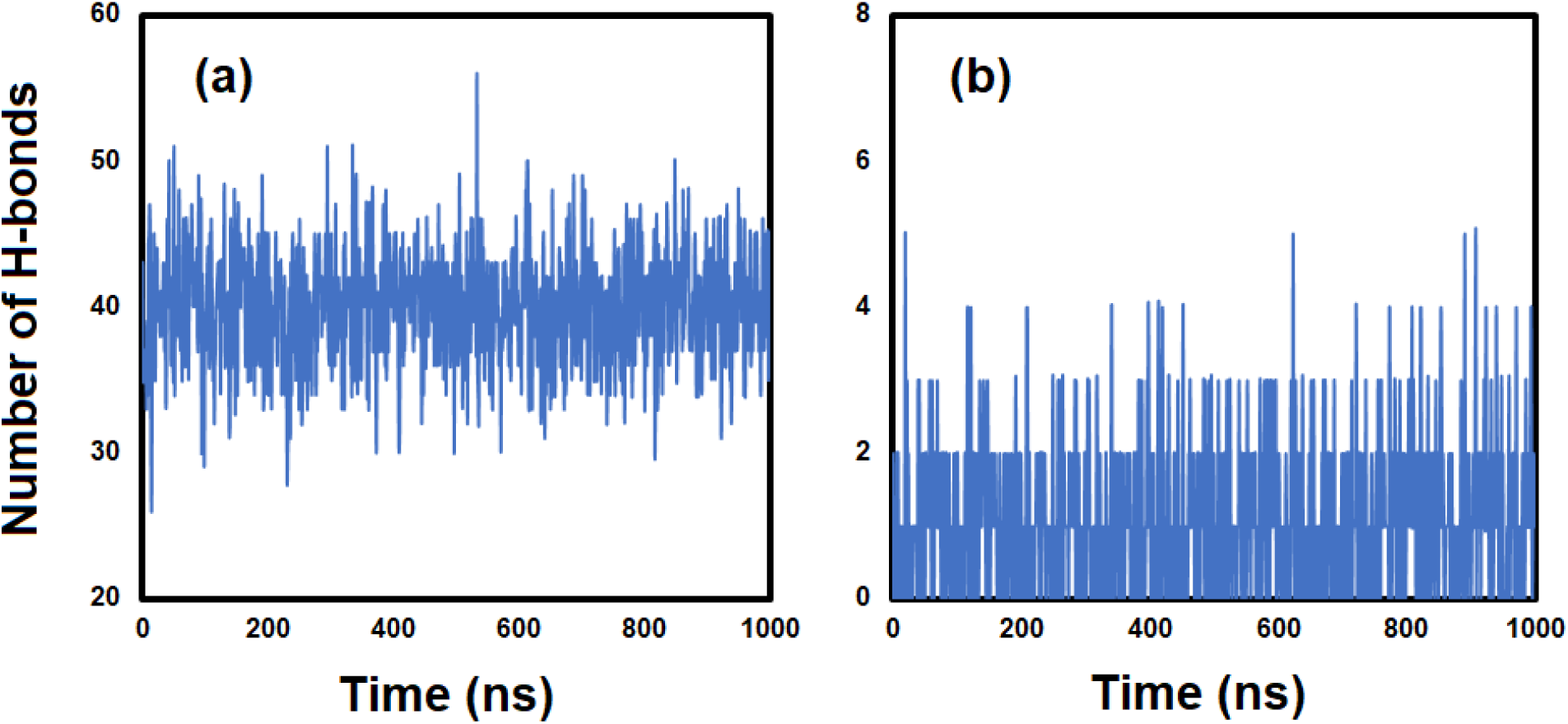
(a) Hydrogen bonds between PHMB and solvent and (b) PHMB intra-molecular hydrogen bonds in 1 PHMB polymer-solvent system.

**Figure S5.**
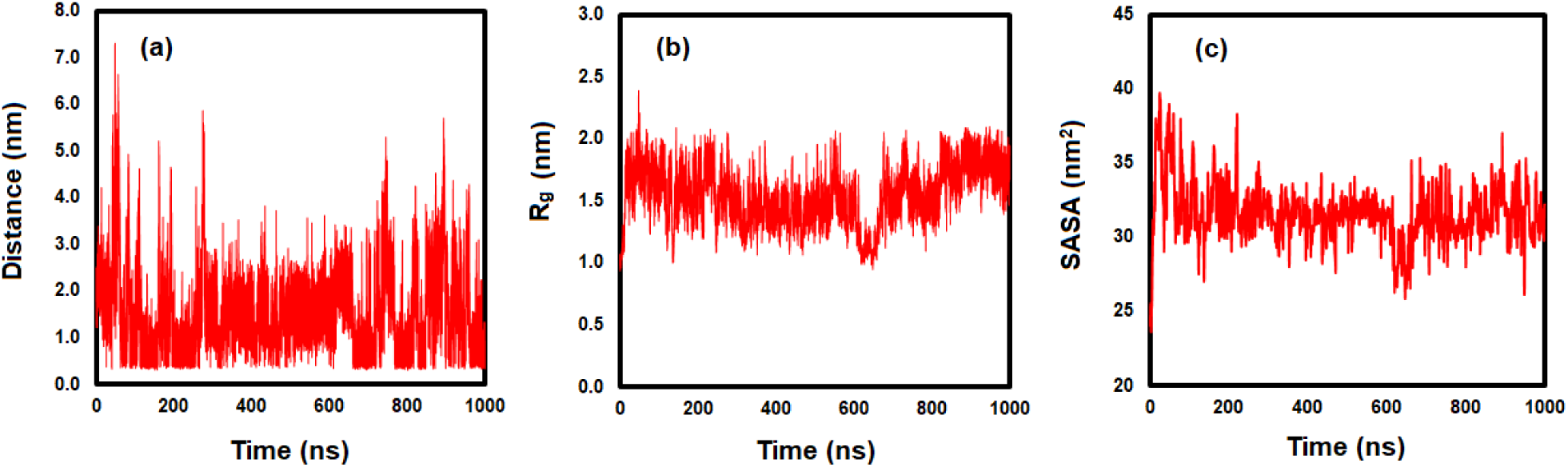
(a) End-to-end distance, (b) radius of gyration and (c) solvent accessible surface area (SASA) of PHMB polymer in solvent. SASA was calculated based on the numerical Double Cubic Lattice Method (DCLM)^1^ as implemented in the GROMACS package using 1 ns intervals.

**Figure S6.**
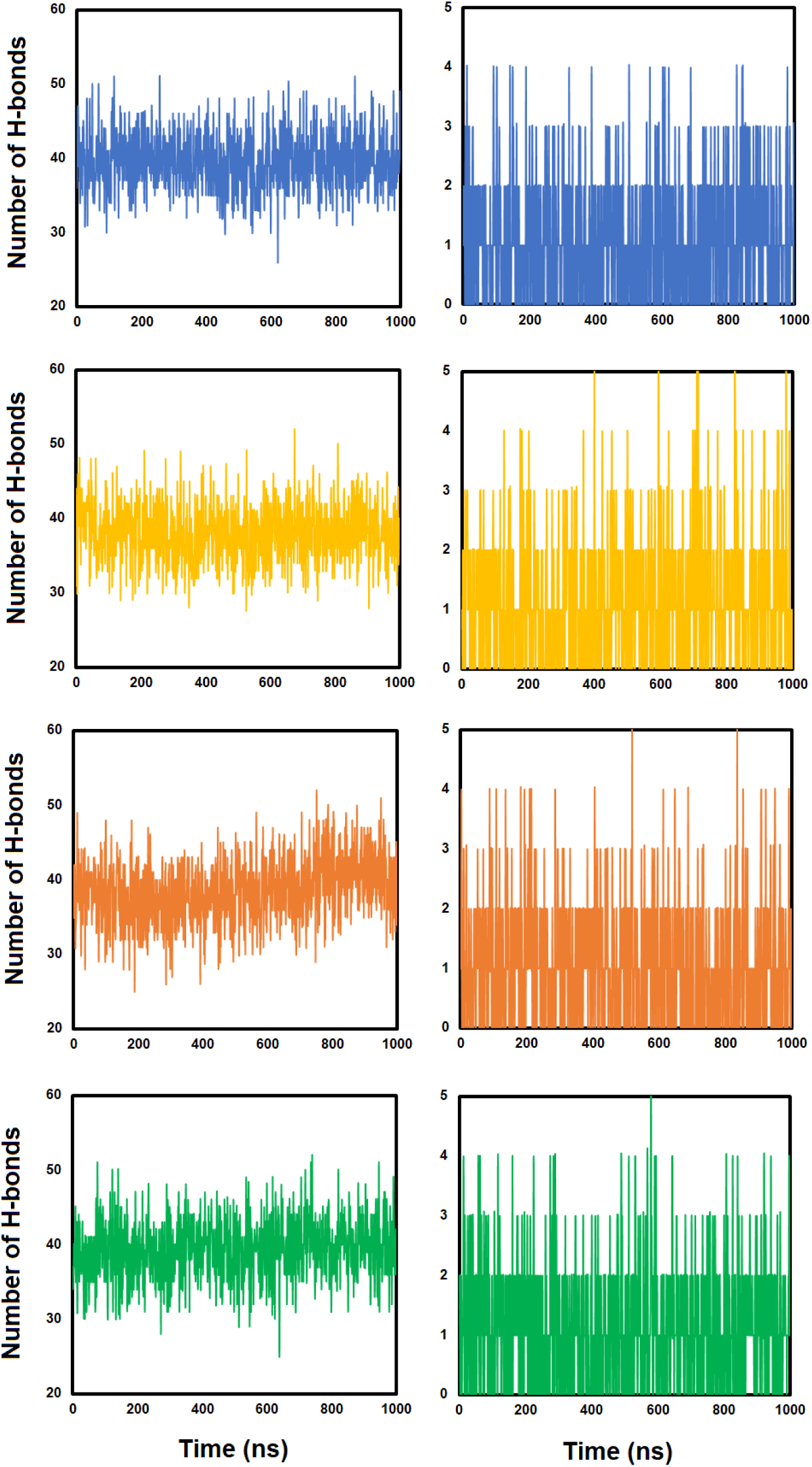
Left panel: Hydrogen bonds between dodecamers and solvent, and right panel: PHMB intra-molecular hydrogen bonds in four dodecamer-solvent system. Each row (shown in different colors) represents one of the dodecamer chains.

**Figure S7.**
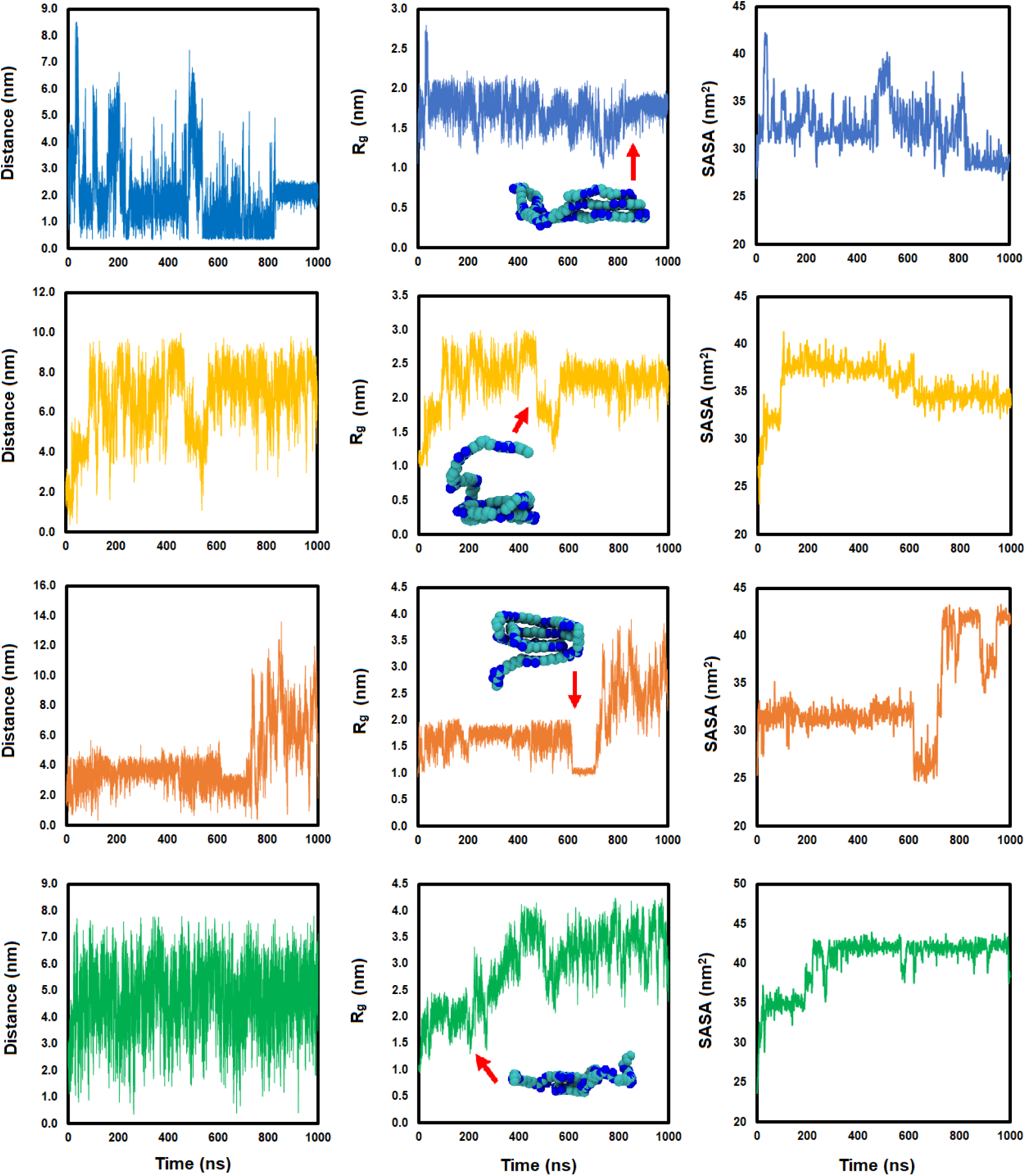
Left panel: End-to-end distance, middle panel: radius of gyration and right panel: solvent accessible surface area (SASA) of each individual PHMB dodecamer molecule in the four dodecamer-solvent system. Each row (shown in different colors) represents one of the dodecamer chains. SASA was calculated based on the Double Cubic Lattice Method (DCLM)^1^ as implemented in the GROMACS package using 1 ns intervals.

**Figure S8.**
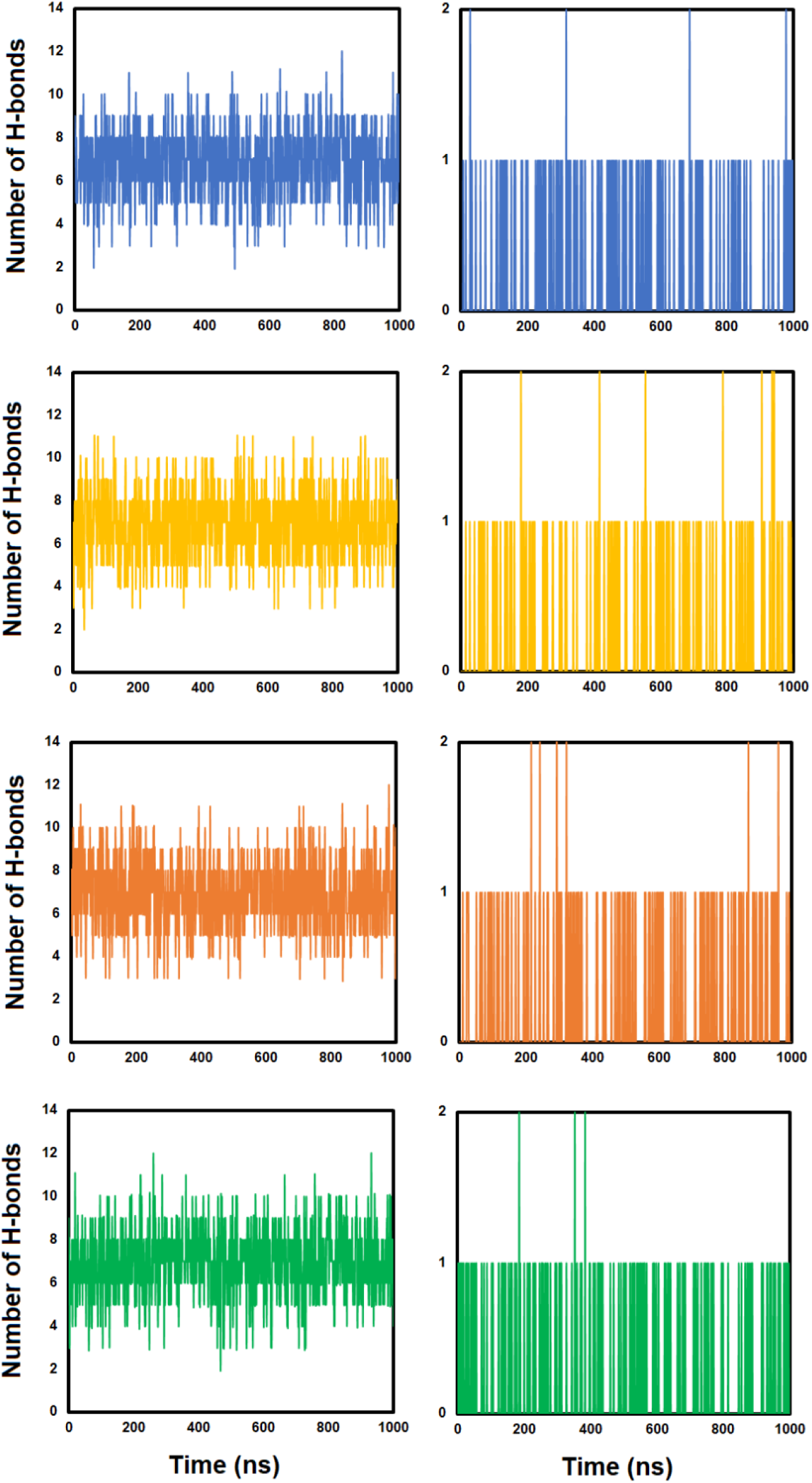
Left panel: Hydrogen bonds between PHMB dimers and solvent. Right panel: PHMB intra-molecular hydrogen bonds in four PHMB dimers-solvent system. Each row (shown in different colors) represents one of the dimer chains.

**Figure S9.**
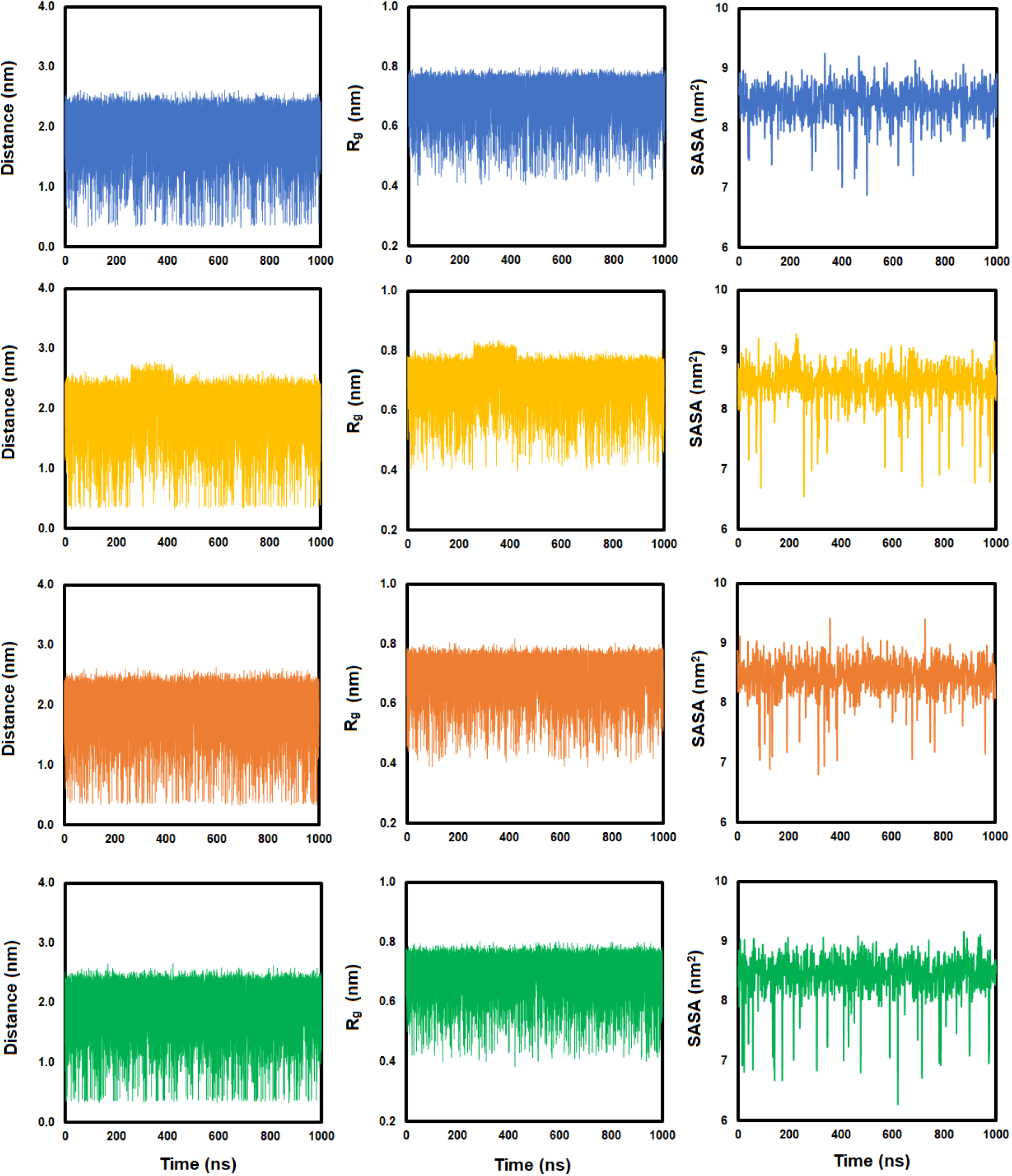
Left panel: End-to-end distance, middle panel: radius of gyration, and right panel: solvent accessible surface area (SASA) of each individual PHMB dimer in the four dimer-solvent system. Each row (shown in different colors) represents one of the dimer chains. SASA was calculated based on the Double Cubic Lattice Method (DCLM)^1^ as implemented in the GROMACS package using 1 ns intervals.

**Figure S10.**
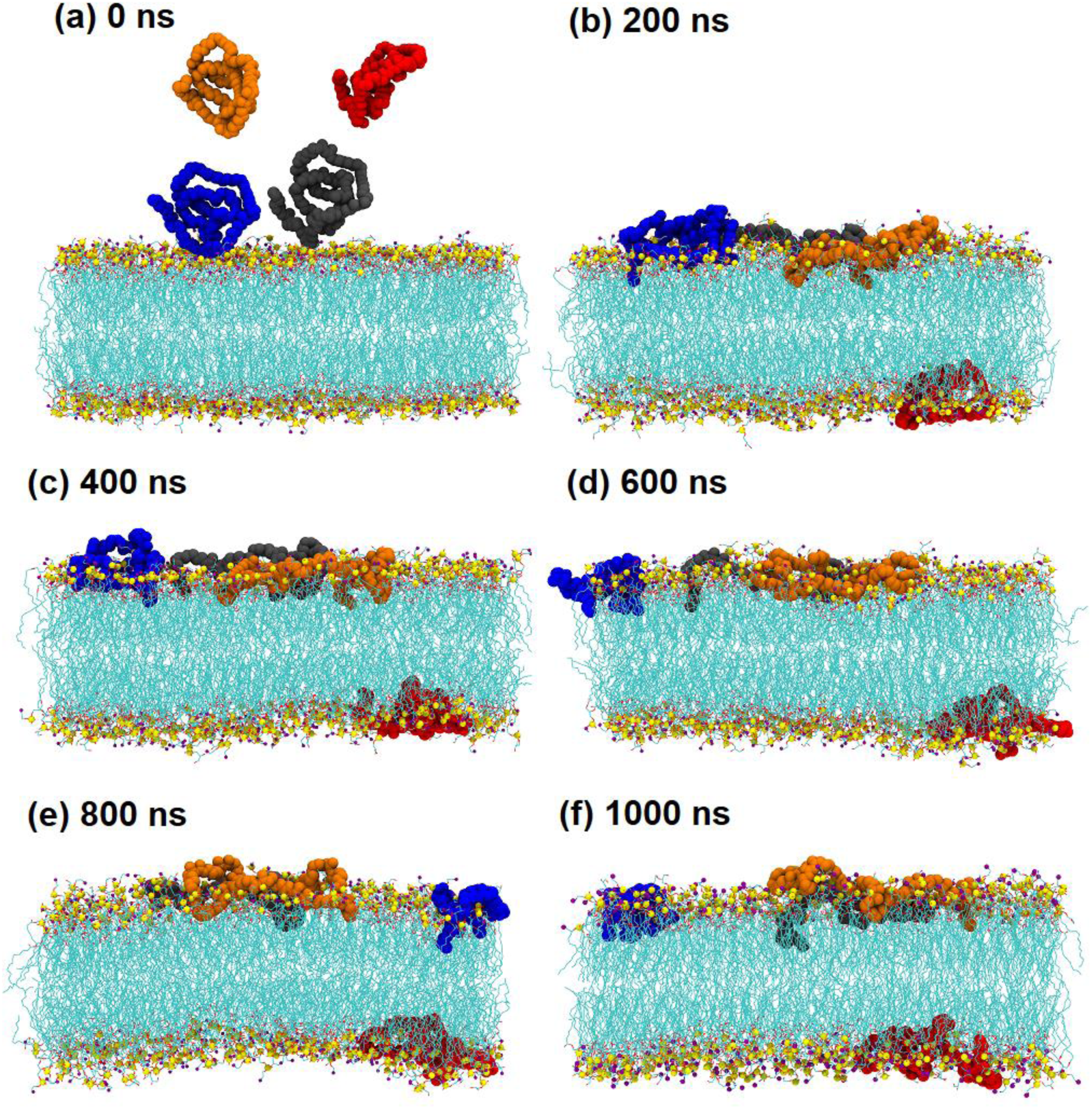
Snapshots of the four PHMB polymers at different times. Nitrogen atoms are shown in purple, phosphorus in yellow and oxygen in red.

**Figure S11.**
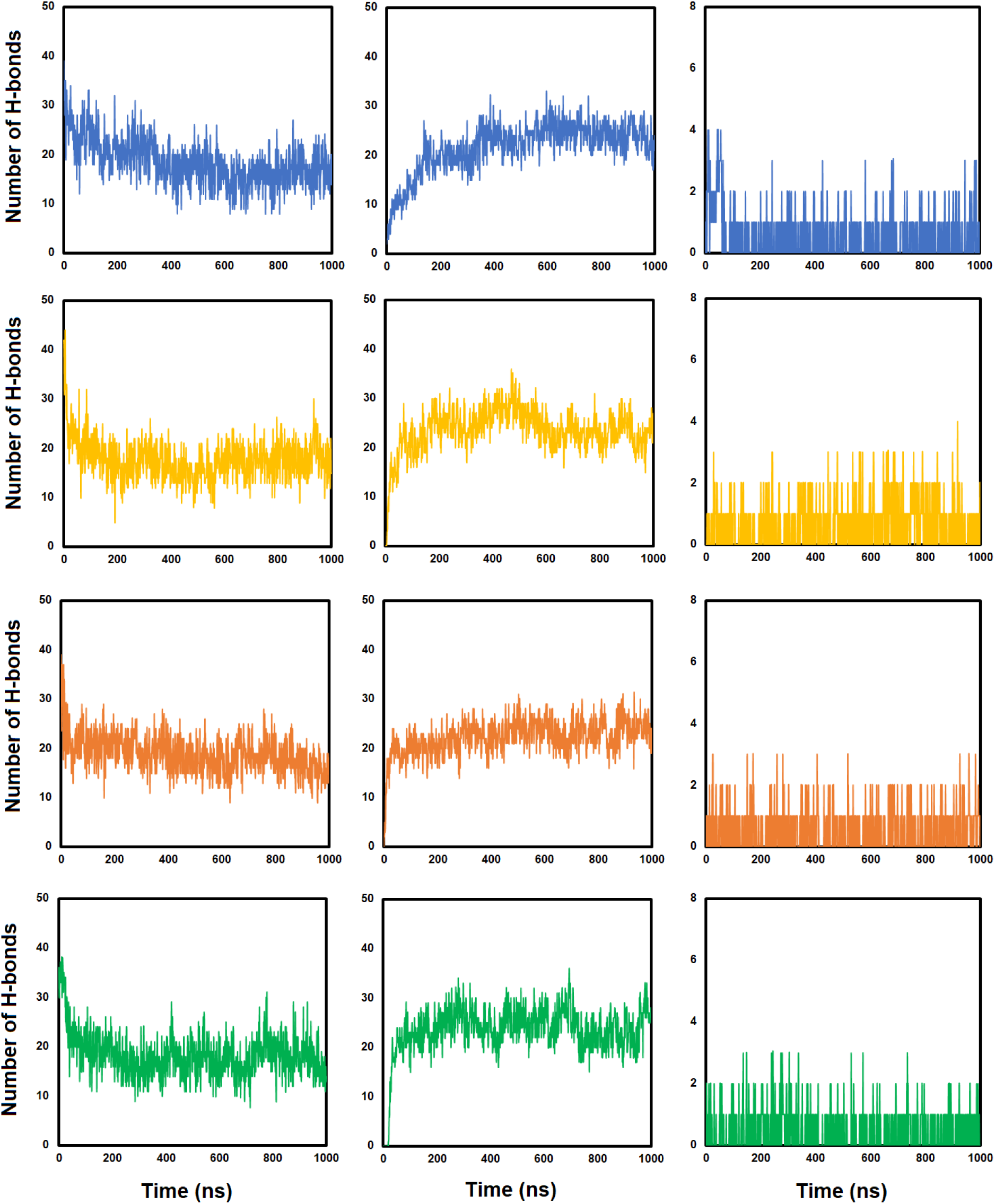
Left panel: Hydrogen bonds between PHMB dodecamers and solvent. Middle panel: PHMB dodecamer-bilayer hydrogen bonds, and right panel: PHMB intra-molecular hydrogen bonds in four PHMB dodecamer-solvent system. Each row (shown in different colors) represents one of the dodecamer chains.

**Figure S12.**
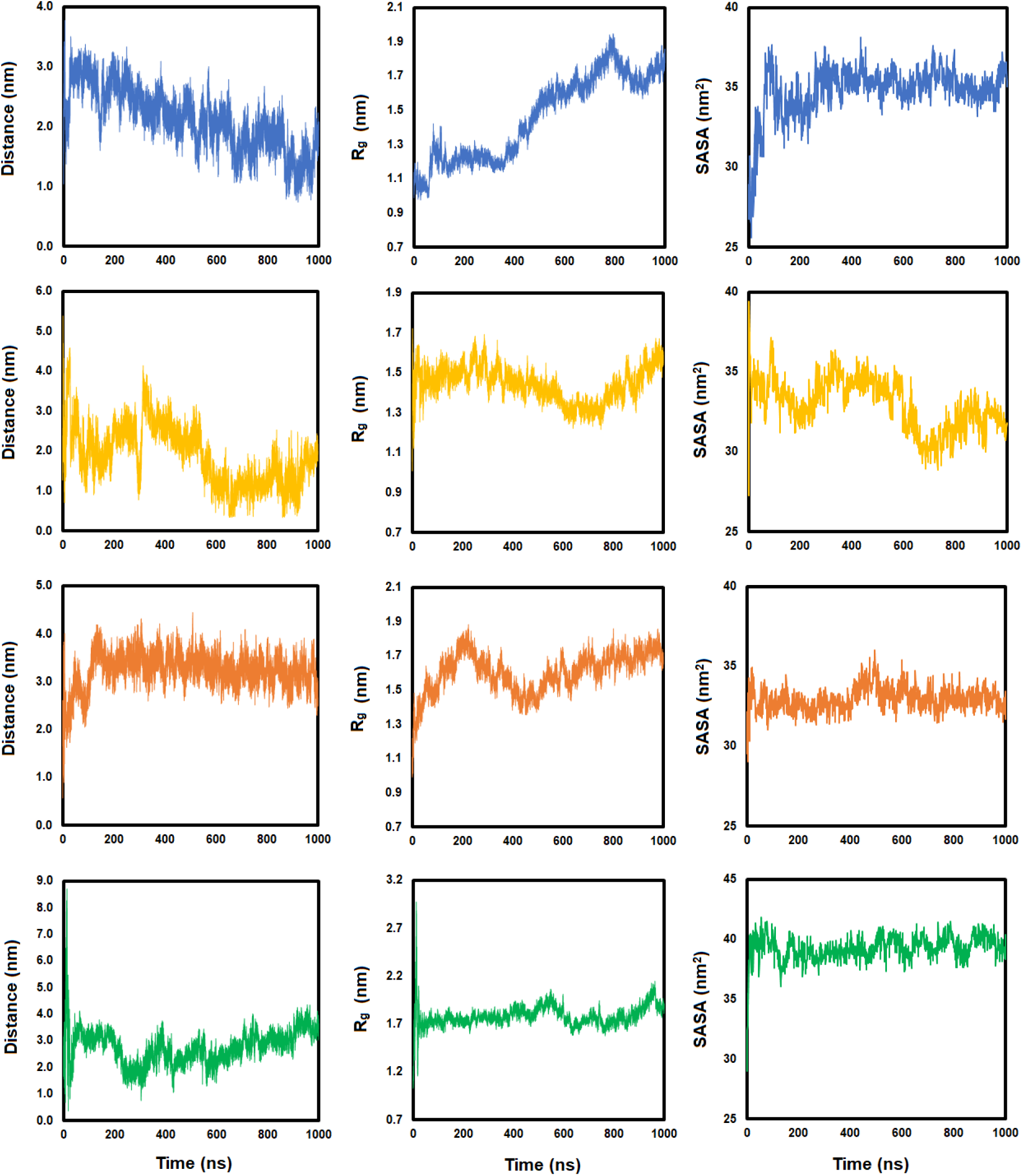
Left panel: End-to-end distance, middle panel: radius of gyration and right panel: solvent accessible surface area (SASA) of each individual PHMB dodecamer in the four dodecamer-bilayer system. Each row represents one of the dodecamer chains. SASA was calculated based on the Double Cubic Lattice Method (DCLM)^1^ as implemented in the GROMACS package using 1 ns intervals.

**Figure S13.**
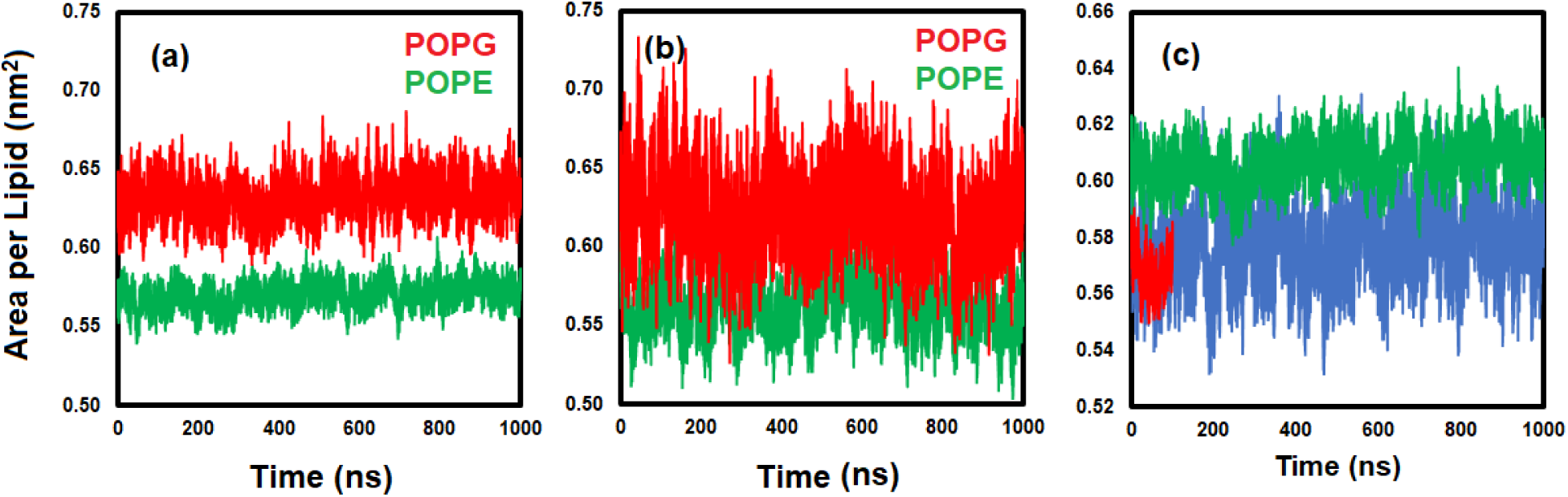
Area per lipid (red: POPG, green: POPE) over time for (a) four PHMB polymers-bilayer and (d) four PHMB dimers-bilayer systems. (c) Overall area per lipid for (red) pure bilayer, (green) four PHMB dodecamers-bilayer, and (blue) four PHMB dimers-bilayer.

**Figure S14.**
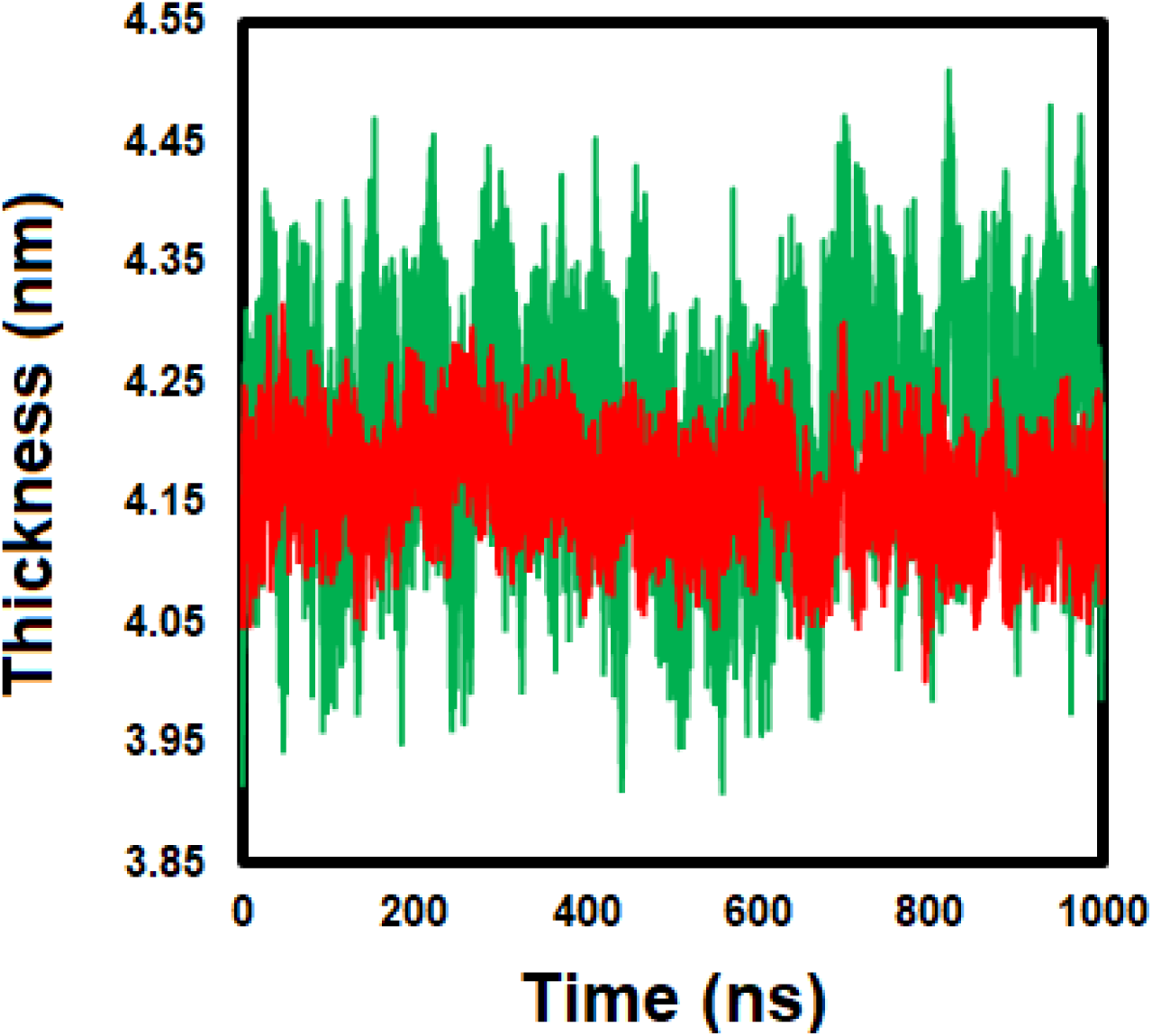
Bilayer thickness over time for (red) four PHMB dodecamers-bilayer and (green) four PHMB dimers-bilayer systems based on the average position of the PO_4_ groups (see Figure S2 for the structure).

**Figure S15.**
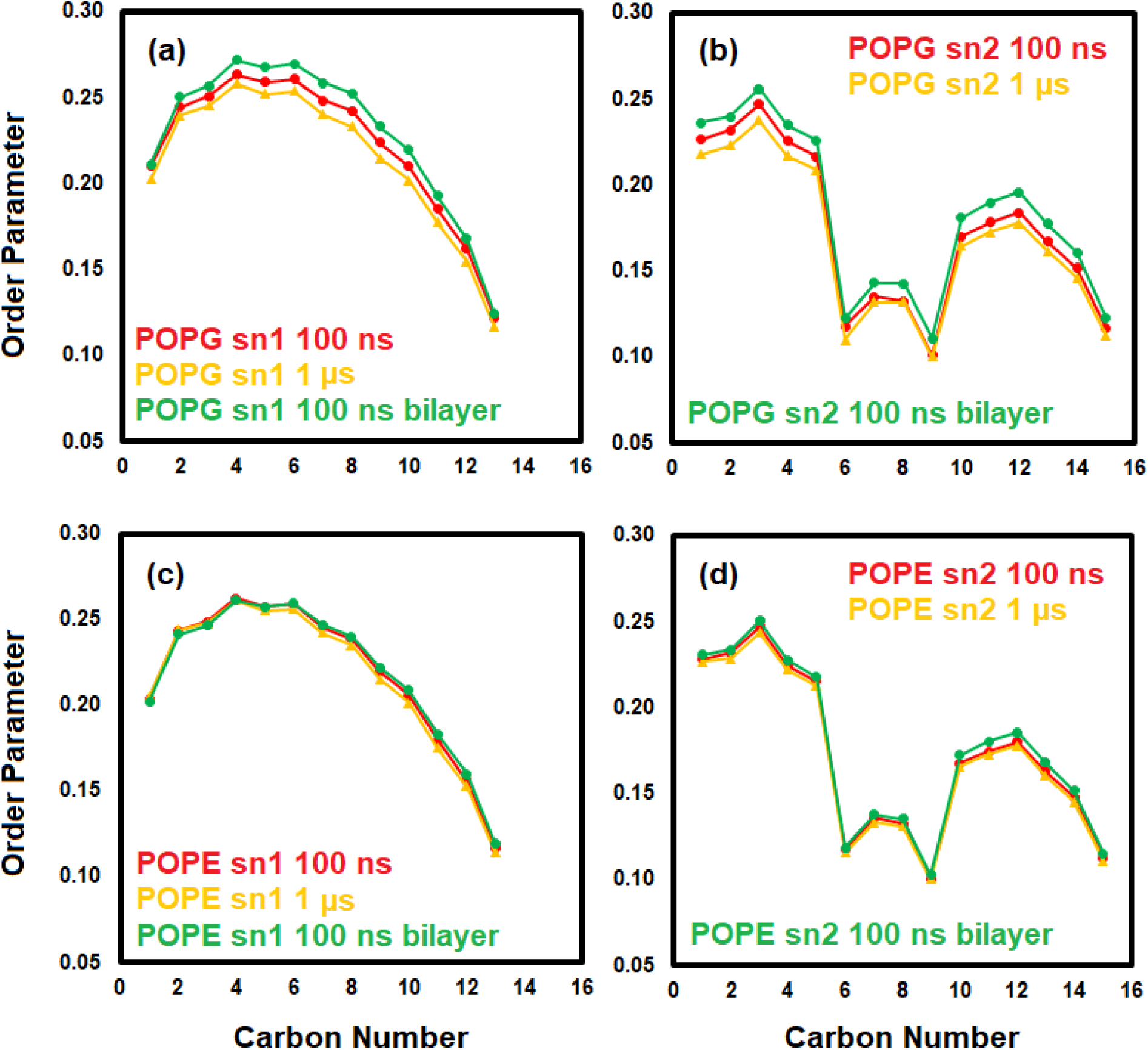
Order parameter of (a) POPG sn-1, (b) POPG sn-2, (c) POPE sn-1, and (d) POPE sn-2 chains (0 – 100 ns) (green for the bilayer in the absence of the polymer) and (900 – 1000 ns) for the 4 PHMB dodecamers-bilayer system.

**Figure S16.**
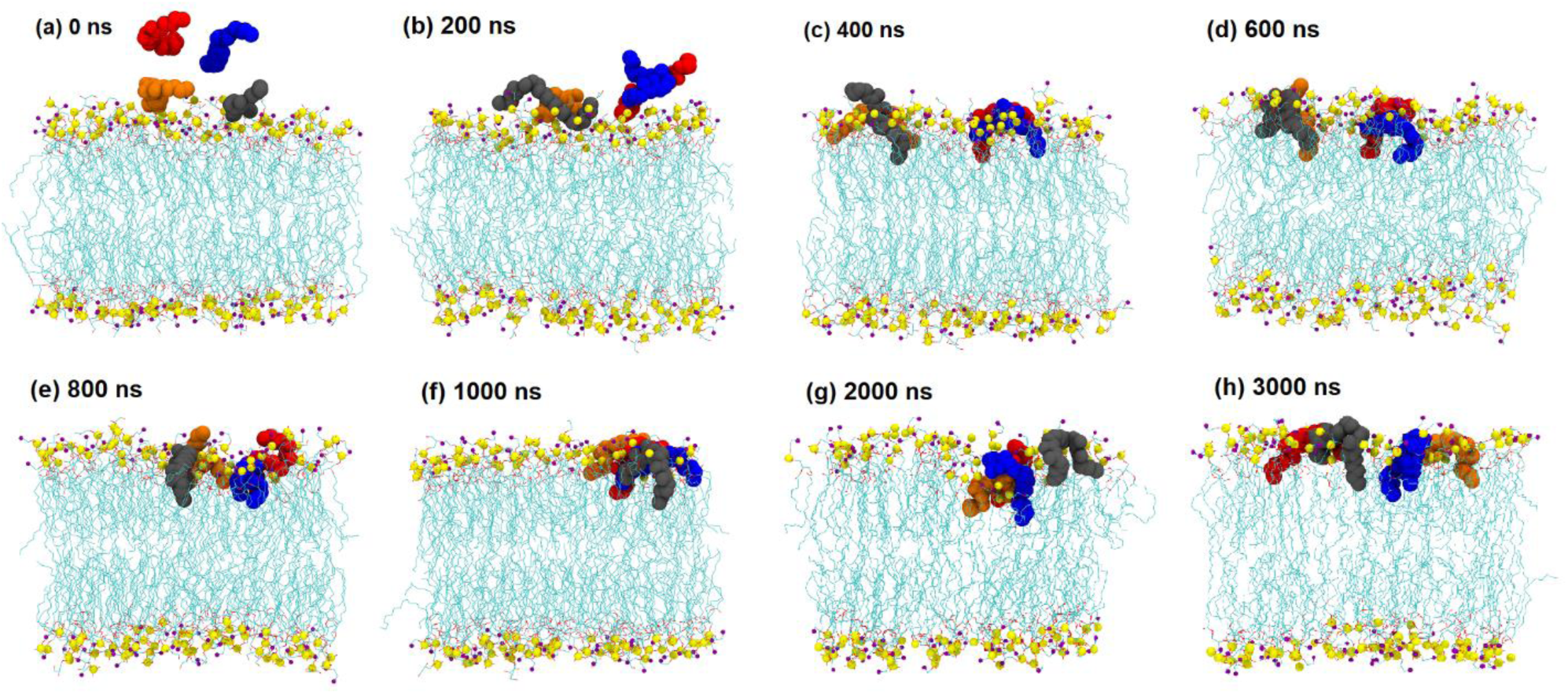
Snapshots of the four at different times. Nitrogen is shown in purple, phosphorus in yellow, and oxygen in red.

**Figure S17.**
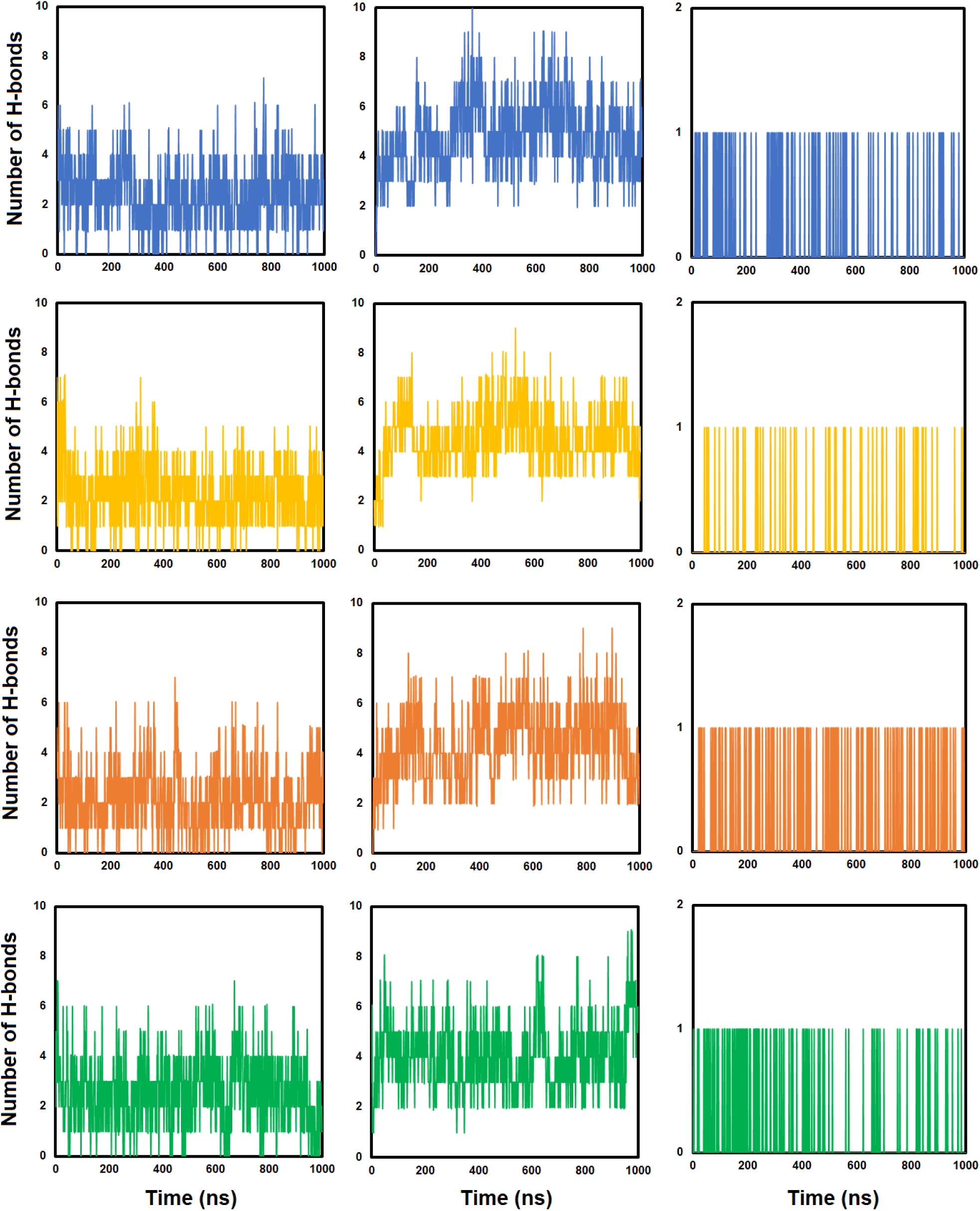
Left panel: Hydrogen bonds between dimers and solvent, middle panel: PHMB dimer-bilayer hydrogen bonds, and right panel: PHMB intra-molecular hydrogen bonds in four dimer-solvent system. Each row (shown in different colors) represents one of the dimer chains.

**Figure S18.**
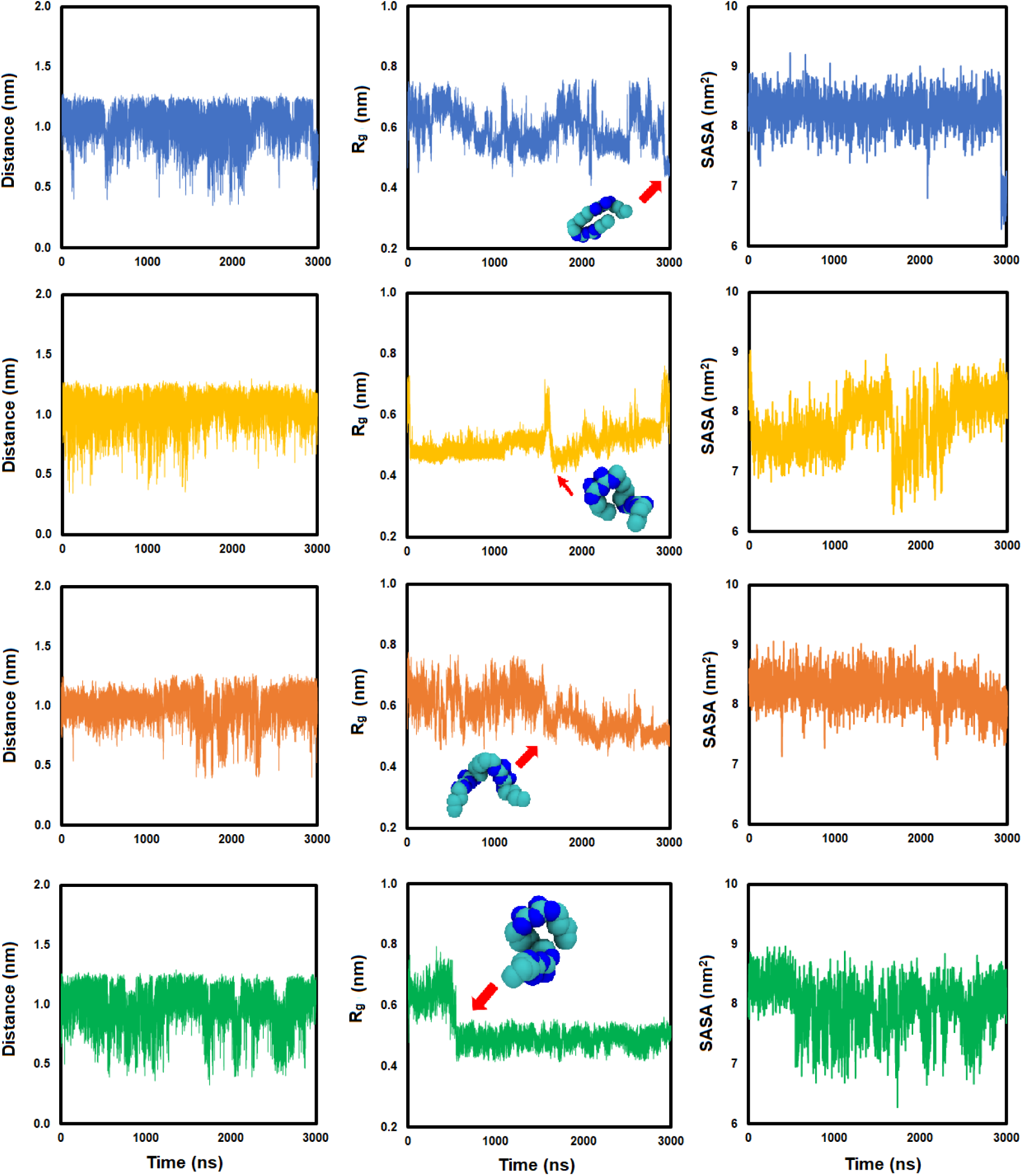
Left panel: End-to-end distance, middle panel: radius of gyration, and right panel: solvent accessible surface area (SASA) of each individual PHMB dimer in the four dimer-bilayer system. Each row (shown in different colors) represents one of the dimer chains. SASA was calculated based on the Double Cubic Lattice Method (DCLM) as implemented in the GROMACS package using 1 ns intervals.

**Figure S19.**
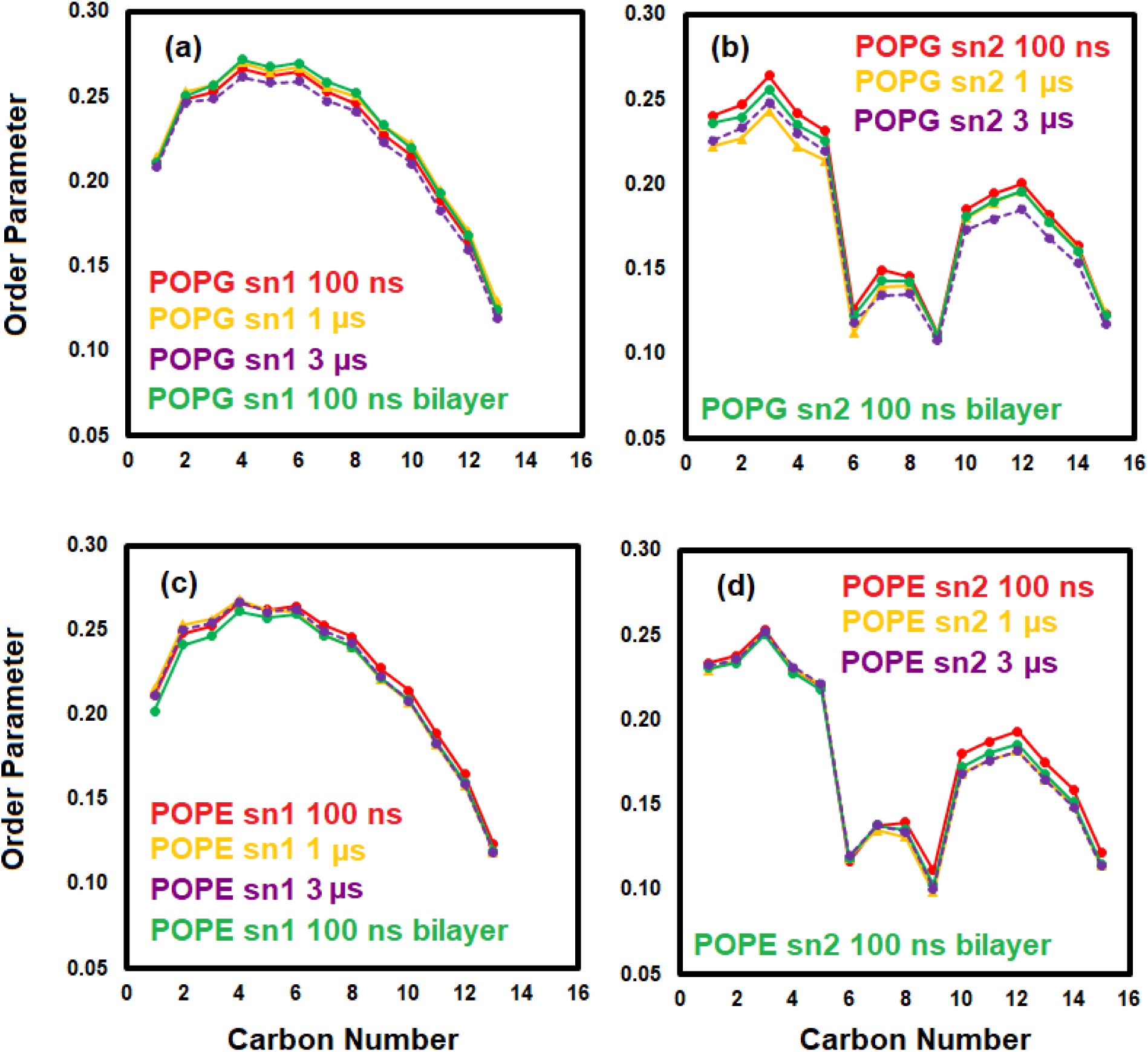
Order parameter of (a) POPG sn-1, (b) POPG sn-2, (c) POPE sn-1, and (d) POPE sn-2 chains (0 – 100 ns) (green for bilayer in the absence of the PHMB chains) and (900 – 1000 ns) (orange) and (2900 – 3000 ns dashed purple) for the four PHMB dimers-bilayer system.

**Figure S20.**
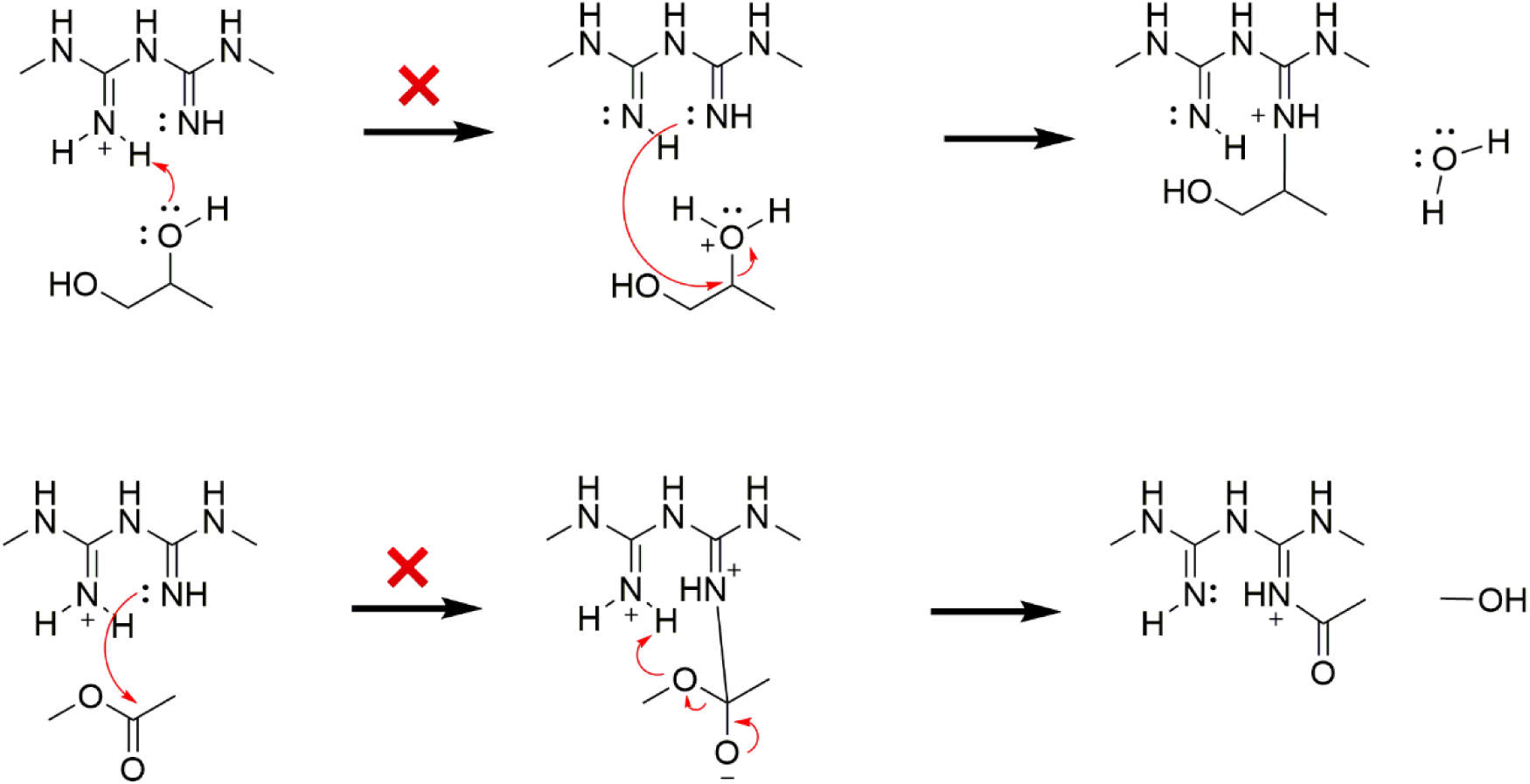
Potential reactions between biguanide and phospholipids that could not be characterized. Transition structures for these reactions were not found due to the unlikeliness of the reactions.

**Figure S21.**
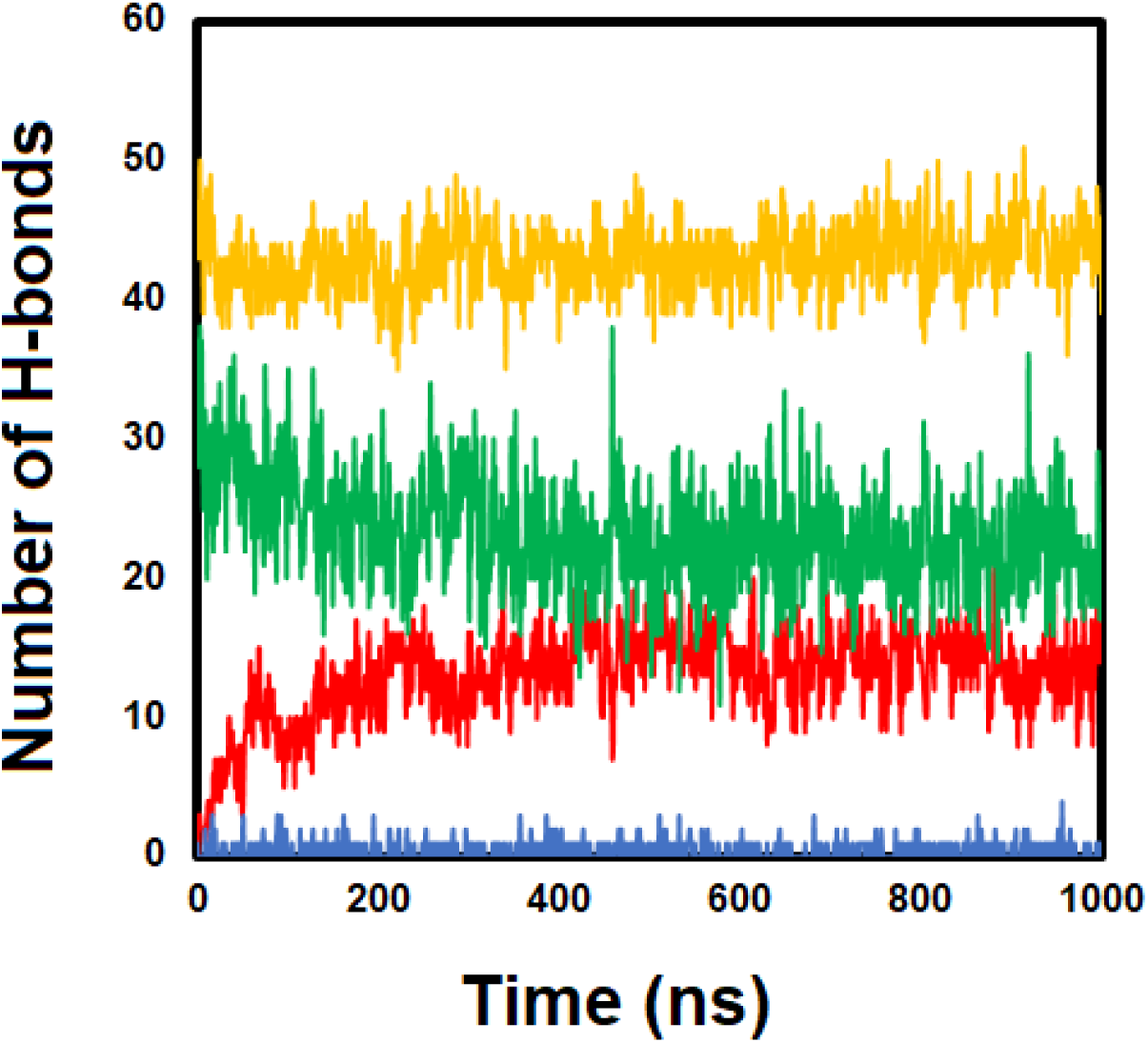
Average number of base pair hydrogen bonds in DNA (orange), PHMB–DNA (red), PHMB–solvent (green), and PHMB-PHMB (blue) hydrogen bonds.

**Figure S22.**
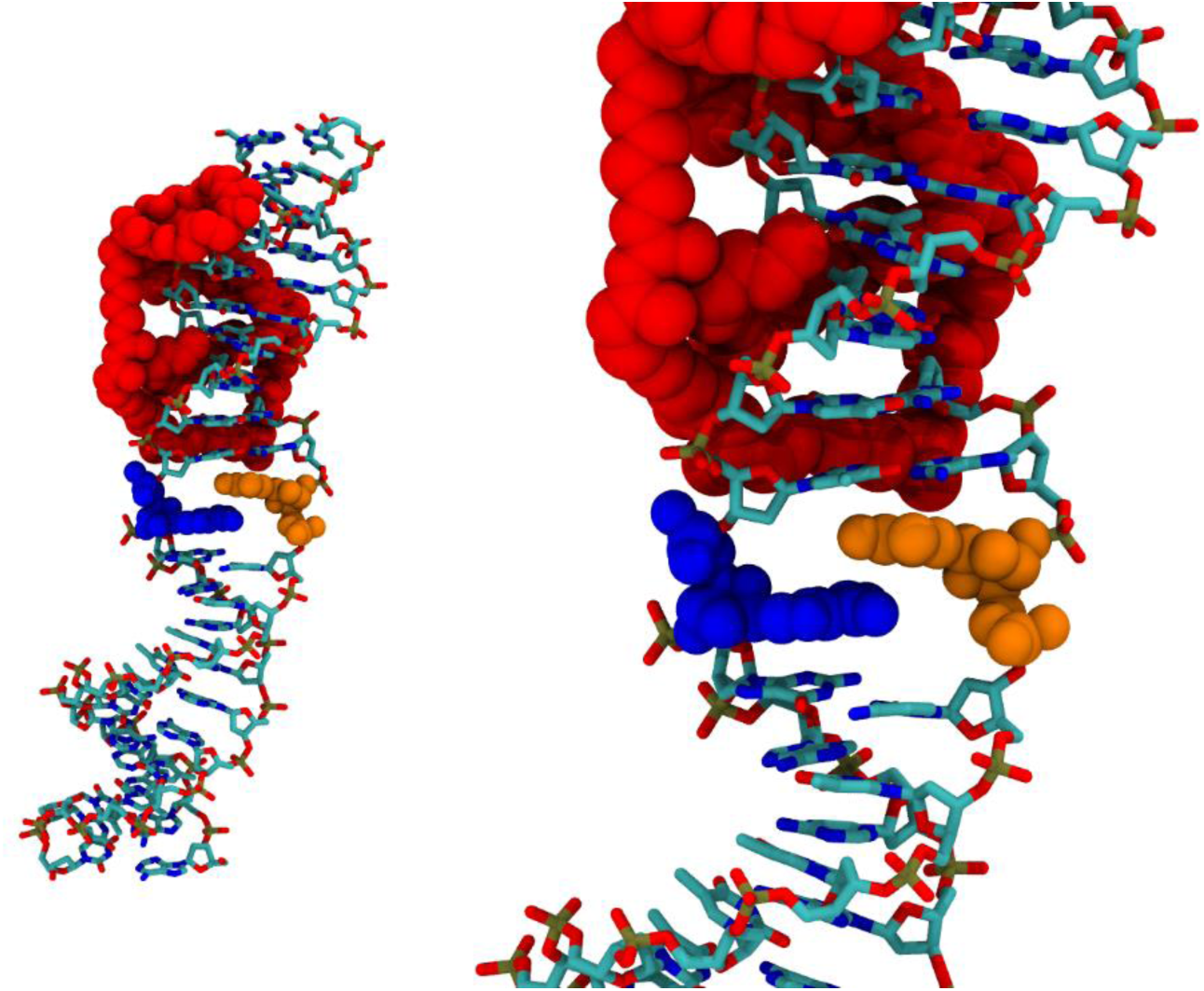
Base-pair dislocation in the DNA oligomer and its relative position to PHMB polymer.

**Figure S23.**
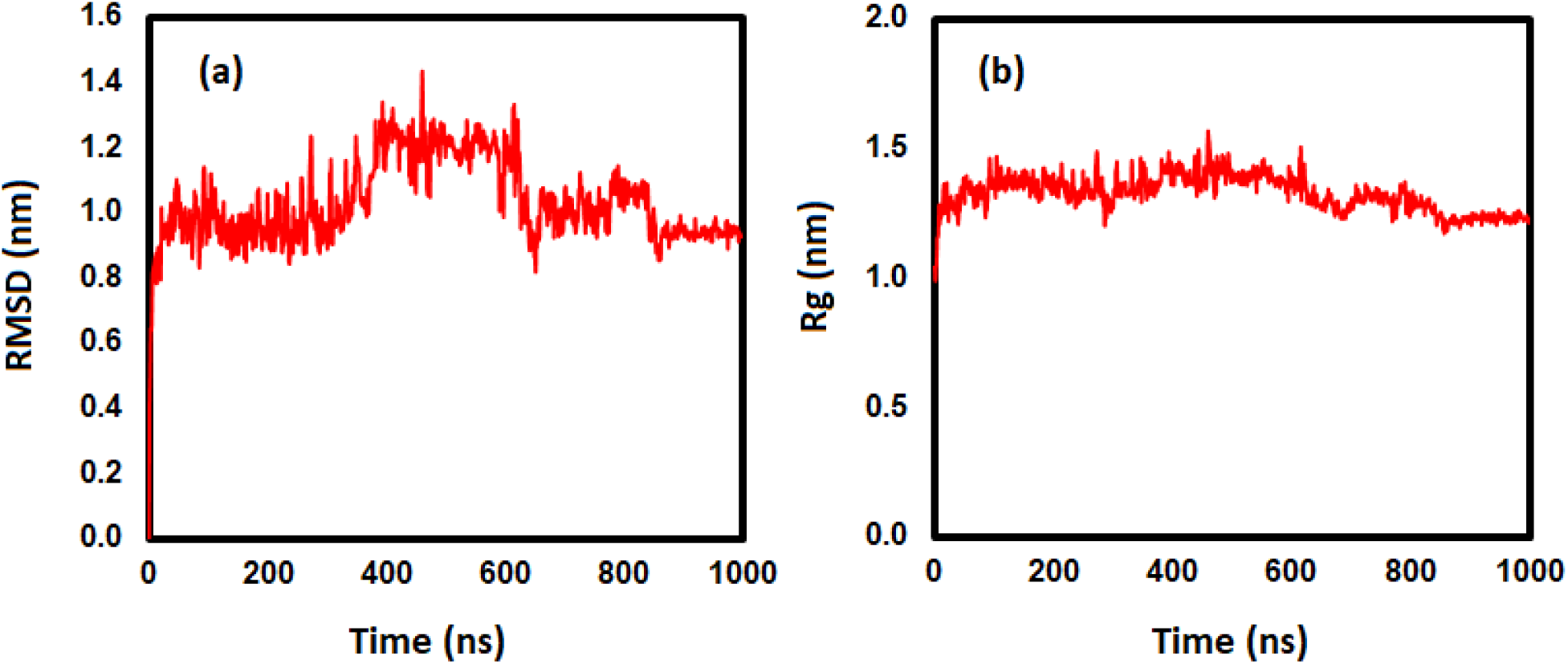
(a) RMSD and (b) radius of gyration of PHMB in the polymer–DNA system.

